# Integration of clinical and T-cell immune profiling to predict early response to CD3xBCMA bispecific antibody immunotherapy in Multiple Myeloma

**DOI:** 10.64898/2026.08.17.743749

**Authors:** Nicolas Deredec, Lisa Aziez, Ismael Boussaid, Justine Decroocq, Axelle Guedon, Margaux Michot, Cyril Catelain, Dorothée Selimoglu-Buet, Ahmadreza Arbab, Cécile Alanio, Olivier Kosmider, Lise Willems, Michaela Fontenay, Patricia Franchi, Rudy Birsen, Nicolas Chapuis, Didier Bouscary, Marguerite Vignon, Yannick Simoni

## Abstract

The emergence of bispecific antibodies (BsAbs) targeting T cells (CD3^+^) and tumor plasma B cells (BCMA^+^) has provided a new therapeutic option for patients with relapsed/refractory multiple myeloma cancer. However, responses to CD3×BCMA BsAb therapy remain heterogeneous, and treatment is associated with frequent immune-related adverse events. Although baseline immune characteristics have been associated with clinical outcomes, little is known about the early immune dynamics induced by this therapy. Here, we investigated whether longitudinal clinical monitoring and high-dimensional profiling of blood circulating T cells could identify early biomarkers of response or toxicity during treatment. Our results indicate that all treated patients exhibit an early depletion of circulating T cells associated with T-cell activation within the first two weeks. Integration of clinical and immunological parameters using Factorial Analysis of Mixed Data (FAMD) identified immune features associated with treatment outcome. Responders had lower plasma soluble BCMA concentrations, fewer bone lesions, higher circulating lymphocyte counts at baseline. During the first days of treatment, responders exhibited a more pronounced increase in plasma CXCL10 levels, associated with a greater decrease in T lymphocyte counts. Overall, our findings suggest that integrating clinical and immune parameters measured during the first days of treatment may enable early patient stratification and support the development of a predictive score to identify patients with multiple myeloma who are most likely to benefit from CD3×BCMA BsAb therapy.

**Graphical Abstract:** 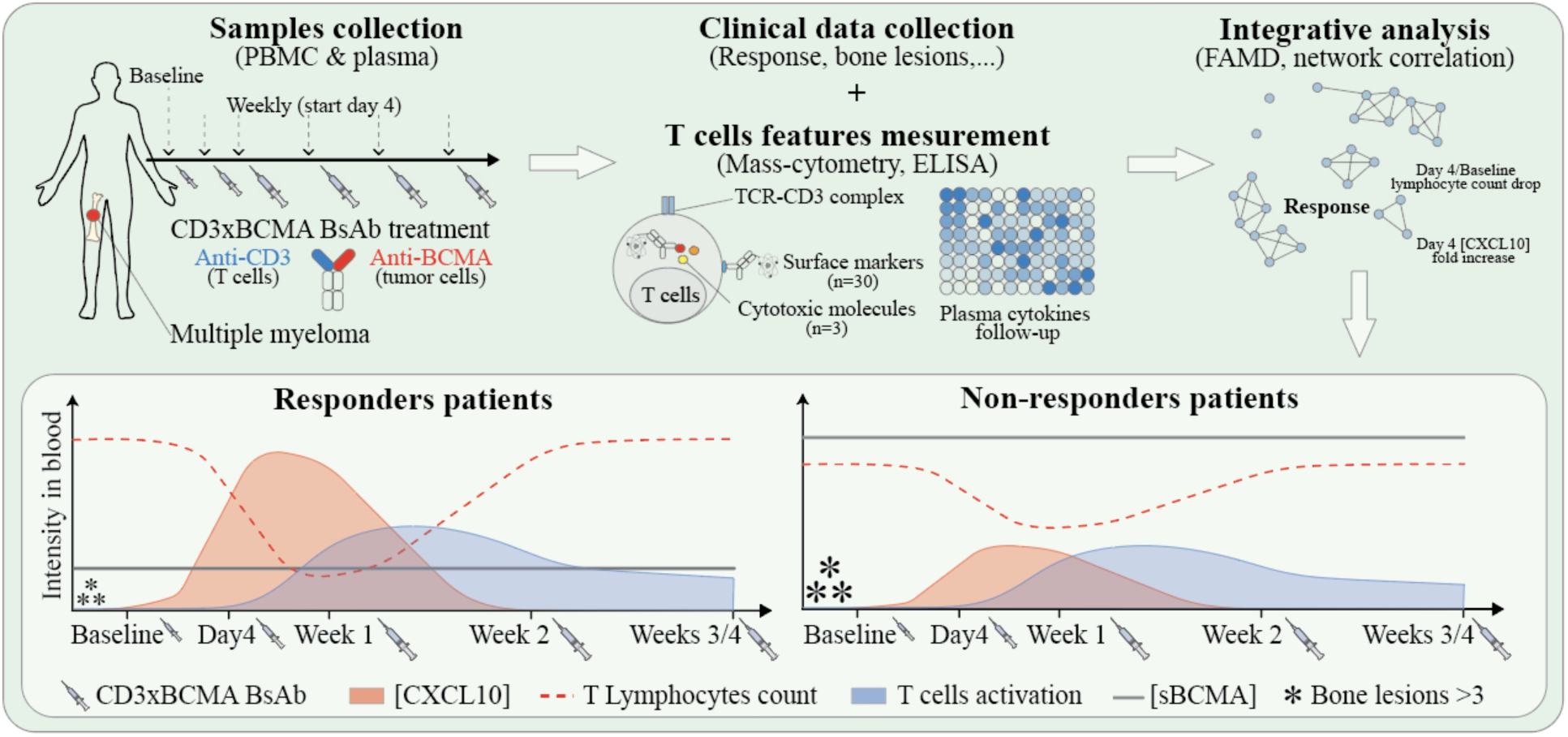

**Highlights:**

- Integrated clinical and blood T-cell immune profiling using FAMD enables patient stratification following CD3xBCMA BsAb therapy.
- T-cell immune activation occurs predominantly within the first two weeks of therapy.
- First-week clinical and immune parameters identify patients most likely to benefit from therapy.
- High CXCL10 levels, a profound early decline in circulating T cells, low sBCMA levels, and fewer bone lesions are candidate predictive markers of treatment response.

## Introduction

Despite decades of research and the approval of 10 different therapeutic classes^1^, multiple myeloma (MM) remains an incurable blood cancer. Characterized by the abnormal proliferation of malignant plasma B cells within the bone marrow, MM accounts for approximately 10% of all hematologic cancers^2,3^. Clinically, the disease is associated with a broad spectrum of complications, including osteolytic bone disease (79%), anemia (73%), and acute kidney injury (19%) among patients^3^. Current treatment strategies combine proteasome inhibitors, immunomodulatory drugs, and anti-CD38 monoclonal antibodies^3^. Although these therapies have markedly improved outcomes, doubling the 5-year survival rate since 1990^4^, most patients ultimately relapse^5^.

The emergence of bispecific antibodies (BsAbs) has recently expanded treatment options for patients with relapsed or refractory MM. Among these agents, teclistamab and elranatamab simultaneously engage CD3 receptor on T cells and B-cell maturation antigen (BCMA) on tumor cells, redirecting cytotoxic T lymphocytes toward malignant plasma B cells (CD3×BCMA BsAbs). Clinical trials of teclistamab and elranatamab have reported incomparable outcomes, with overall response rates of 63.0% and 63.6% and median durations of response of 18.4 and 17.1 months, respectively^6,7^. However, treatment is associated with frequent adverse events, including cytokine release syndrome (CRS), cytopenias, and atypical infections. Furthermore, response to CD3×BCMA BsAb therapy varies substantially between patients. Several determinants of therapeutic efficacy have been identified on both the tumor and immune sides. On the tumor side, high tumor burden and elevated levels of soluble BCMA correlate with poorer outcomes^8,9^. Resistance mechanisms have also been described, including missense mutations or in-frame deletions within the BCMA extracellular domain and rare cases of biallelic loss of TNFRSF17, the gene encoding BCMA^10,11^. On the T cells side, the baseline fitness and composition of the T-cell compartment strongly influence response. Responding patients typically display higher overall T-cell counts and increased frequencies of naïve and effector CD8⁺ T cells^9^. High baseline levels of cytotoxic CD8⁺ T cells and low levels of exhausted CD8⁺ and regulatory T cells have also been associated with improved responses to CD3×BCMA BsAbs therapy^9,12,13,14^. *In vitro* experiments showed that CD3×BCMA BsAb induce T cells activation within 24 h with upregulation of CD69 and CD25, increase their cytotoxic and cytokines releases (IFNγ, TNFα)^15^. However, little is known about how T-cell features evolve during the early phase of this therapy or how dynamic immune parameters interact with clinical variables to influence treatment outcomes. Because these determinants are heterogeneous in nature, spanning clinical features, cytokine measurements, and high-dimensional immunophenotyping, integrative analytical approaches are required to capture their relationships.

In this pilot study, we aimed to identify early determinants of response and toxicity during CD3×BCMA BsAb therapy by integrating longitudinal clinical and immunological data. Blood from multiple myeloma patients was collected at baseline and during the first week of therapy. T-cell lineage, differentiation state and plasmatic cytokines implicated in T-cell activation were measured. These serial heterogeneous datasets composed of immunological and clinical data were subsequently integrated using factorial analysis of mixed data (FAMD) and represented by a correlation network, allowing comprehensive analysis and rapid identification of clinical and immunological parameters correlating with response.

## Results

### CXCL10 induction and transient T-cell lymphopenia within the first week characterize patients treated with CD3×BCMA BsAb therapy

Cytokines and chemokines associated with T-cell activation and recruitment were measured by ELISA in patients’ plasma (n=14) at baseline, before each escalation dose within the first week (i.e. Day 4 and Week 1) and weekly throughout the first month of CD3×BCMA BsAb therapy (Figure 1A, Table S1). High plasma TNFα concentrations were observed at baseline (Mean=151.4pg/mL) compared to healthy donors (HD, Mean=2.9pg/mL). The inter-patient variability suggests pre-existing systemic inflammation in a subset of patients. This variability persisted, with no significant overall changes observed following therapy initiation. IFNγ levels were low in patients at baseline (Mean=11.8 pg/mL) compared with healthy donors (Mean=20.1pg/mL). Longitudinal monitoring revealed heterogeneous kinetics across patients, with some individuals showing sustained reductions toward physiological levels, while others exhibited persistent or fluctuating IFNγ concentrations. In contrast, plasma CXCL10 concentrations, a chemokine implicated in T cells recruitment at inflamed site, is high at baseline (Mean=77.9pg/mL) compared with healthy donors (Mean=9.8pg/mL) and increased rapidly after treatment initiation, with a significant elevation detected as early as day 4 (299.4 pg/ml). While this early induction was consistently observed across patients, its magnitude and duration varied. CXCL10 levels subsequently declined in several patients, starting at week 1, but remained elevated in a subset (Figure 1B, S1A).

**Figure 1.**
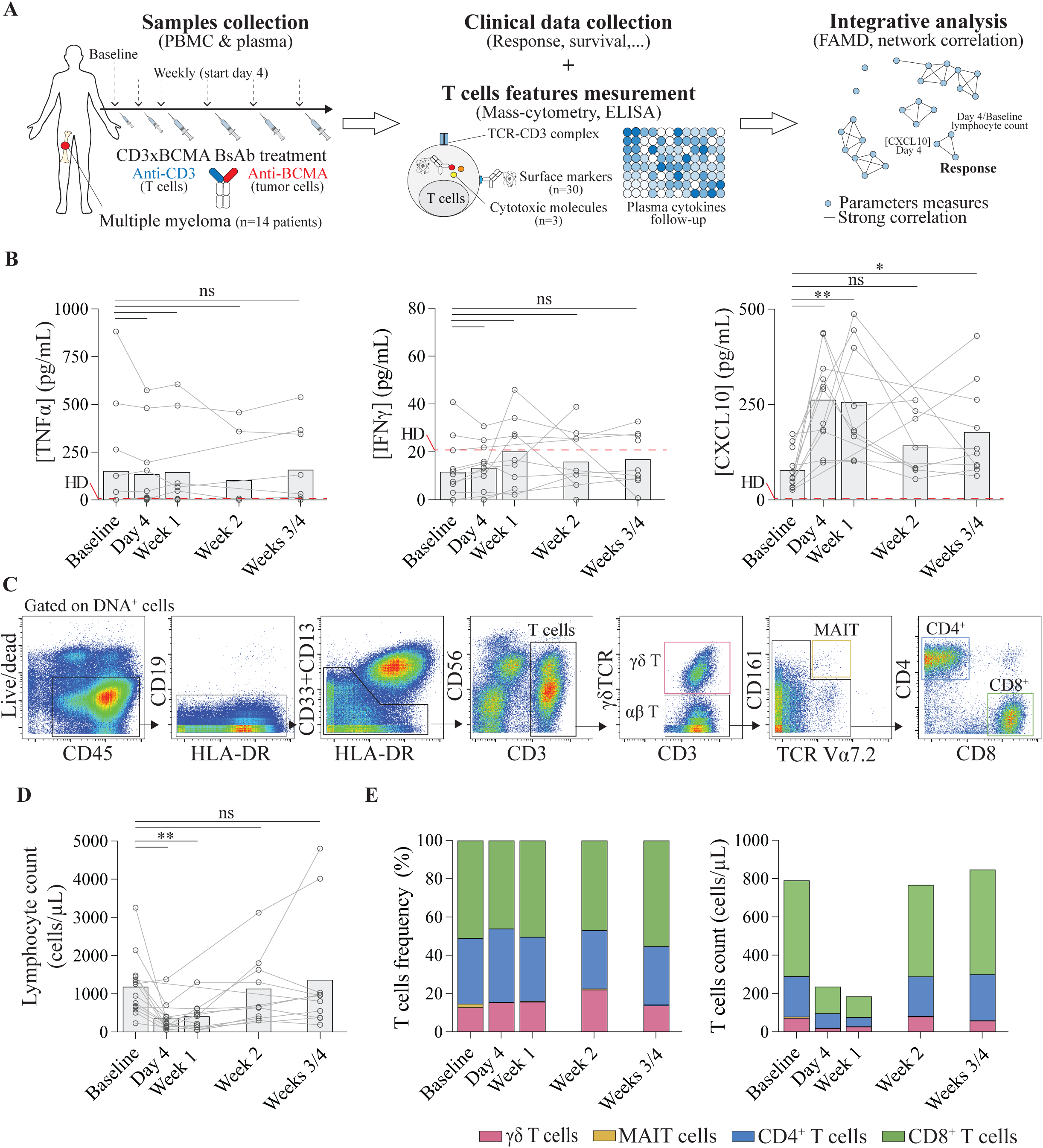
Early T cells changes occur within the first week of CD3xBCMA BsAb therapy. **A.** Schematic overview of sample kinetics and techniques used in this study. **B.** Plasma concentrations of TNFα, IFNγ, and CXCL10 during CD3xBCMA BsAb therapy (n=14 patients). Red dashed lines indicate the average concentration for healthy donors. **C.** Gating strategy used to identify MAIT, CD4^+^, CD8^+^ and γδ T cells from peri- pheral blood mononuclear cells (PBMCs) of patients (gated on DNA^+^ cells). Representative mass cytometry data from one patient. **D.** Circulating lymphocyte count of patients during CD3xBCMA BsAb therapy (n=14 patients) **E.** Frequencies (left panel) and counts (right panel) of MAIT, CD4^+^, CD8^+^ and γδ T cells in patients PBMCs during CD3xBCMA BsAb therapy (n=14 patients). Paired t-tests. ns= non-significant *p ≤ 0.05, **p ≤ 0.01, ***p ≤ 0.001.

To further investigate whether the early increase in plasma CXCL10 was associated with immune cell dynamics, T-cell composition was analyzed by mass cytometry in peripheral blood mononuclear cells (PBMCs) obtained from the same patients and time points as the paired plasma samples (Figure 1A). According to the guidelines for T-cell nomenclature^16^ and using a hierarchical gating strategy, main T-cell lineages were defined for innate γδ T cells and MAIT cells, and conventional CD4⁺ or CD8⁺ T cells (Figure 1C). Longitudinal analysis of patients’ lymphocyte counts revealed a marked and transient decrease at Day 4 following treatment initiation (<200 cells/μL), mirroring the kinetics observed for CXCL10 (Figure 1D). This decrease was already detectable 24 hours after treatment initiation (Figure S1B). Lymphocyte count of patients dropped by 64.1% in the span of 4 days, correlating with their plasma levels of CXCL10 at Day 4 (Figure S1C). Despite this decline, the relative proportions of CD8⁺, CD4⁺, and γδ T cells remained broadly stable throughout treatment, representing approximately 50%, 30%, and 10% of total T cells, respectively. This early and transient lymphopenic phase was followed by a progressive recovery from week 2 onward, with T cell counts returning to baseline levels by weeks 3–4 in most patients (Figure 1E). By contrast, MAIT cells were barely detectable at baseline and remained at very low frequencies throughout treatment in all patients (Figure S1D).

Taken together, these results highlight substantial inter-patient variability at baseline and throughout treatment. Nevertheless, all patients exhibited an early increase in plasma CXCL10 levels accompanied by a transient reduction in circulating T cell counts during the first week of treatment. Importantly, no comparable changes were observed at later time points despite repeated weekly administrations of the CD3×BCMA BsAb.

### Early phenotypic changes of T cells following CD3×BCMA BsAb therapy

To determine whether the transient decrease and subsequent recovery of T cell counts during the first week of treatment were associated with T cell activation, we assessed phenotypic remodeling using UMAP on mass cytometry data. At baseline, UMAP analysis clearly delineated γδ, CD4⁺, and CD8⁺ T cell lineage across patients. However, within each lineage, clustering was strongly driven by patient identity, highlighting substantial inter-patient phenotypic heterogeneity (Figure 2A, 2B, and S2). To account for this variability, analyses were performed on a per-patient basis. Longitudinal UMAP projections revealed dynamic changes in T-cell phenotypic distributions over time. While some patients exhibited minimal phenotypic shifts throughout treatment, others showed early (Day 4) or late (Weeks 3/4) marked redistribution of T-cell populations (Figure 2C). To quantify these heterogeneous dynamics while accounting for inter-patient variability, we compared UMAP distributions at each time point to baseline within each patient using Jensen– Shannon divergence (JS div). This approach enables robust longitudinal comparisons independent of baseline differences, where low JS div (∼0) indicates minimal phenotypic change and high JS div (∼1) reflects strong phenotypic changes from baseline (Figure 2D). Quantitative analysis revealed patient-specific remodeling dynamics emerging as early as Day 4 post-treatment. γδ T cells and CD4⁺ T cells showed a gradual increase in divergence, peaking at Week 2 (∼0.2–0.5), with considerable variability in magnitude and kinetics across patients. CD8⁺ T cells displayed the most pronounced remodeling, with early increases (Day 4) that remained sustained over time, including individuals with strong early shifts (Figure 2E).

**Figure 2.**
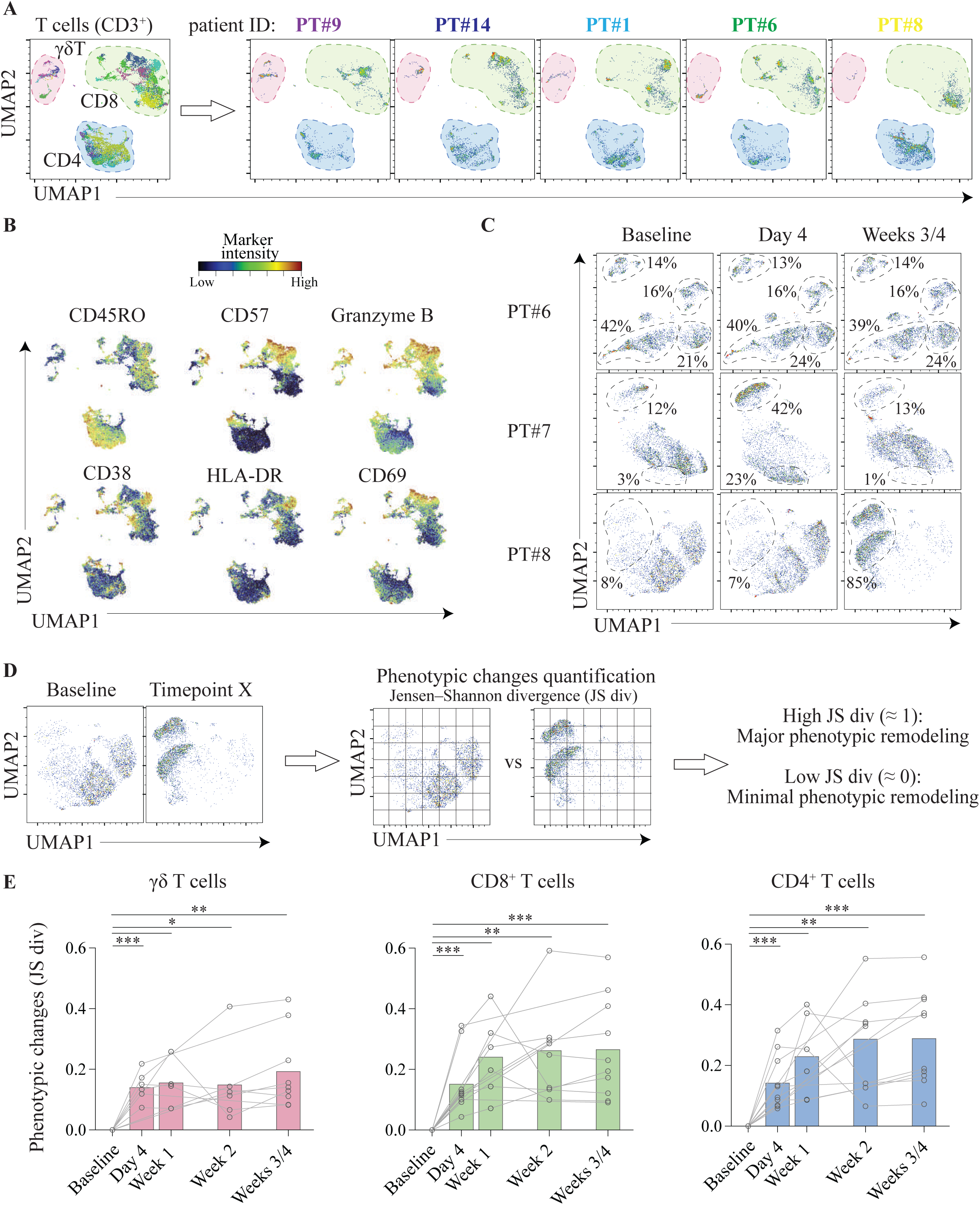
Early phenotypic remodeling of T cells during CD3xBCMA BsAb therapy. **A.** Uniform Manifold Approximation and Projection (UMAP) analysis of CD3⁺ T cells from patients PBMCs at baseline. Combined view (left panel) colored by patient and individual projections (right panel) for 5 patients. **B.** Normalized marker expression intensities were calculated and overlaid on the UMAP plot. **C.** Longitudinal UMAP projections of patients CD8^+^ T cells during CD3xBCMA BsAb therapy. **D.** Schematic representation of the quantification of phenotypic changes by Jensen-Shannon divergence (JS div). **E.** Phenotypic changes (JS div) of γδ, CD4^+^ and CD8^+^ T cells during CD3xBCMA BsAb therapy. Paired t- tests. ns= non-significant *p ≤ 0.05, **p ≤ 0.01, ***p ≤ 0.001.

Collectively, these results indicate that CD3×BCMA BsAb therapy induces early T-cell phenotypic changes during the first week of treatment that vary substantially between patients, reflecting heterogeneous immune responses to therapy.

### Rapid but transient activation of γδ and CD8⁺ T cells during the first two weeks of treatment

To comprehensively analyze phenotypic changes observed using the JS div quantification, we performed manual gating of key activation and differentiation markers across γδ and CD8⁺ T cell lineages. At baseline, with marked inter-patient heterogeneity, innate γδ T cells predominantly displayed a senescent phenotype (CD57⁺ and KLRG1⁺), frequent expression of NK-associated receptors (NKG2C, KIR3DL1, KIR2DL1, and CD226), and strong cytotoxic potential (Perforin, Granzyme A, and Granzyme B) (Figures 3A-C and S3). Following treatment initiation, absolute γδ T-cell counts in peripheral blood decreased during the first week before returning to baseline by Week 2. Apart from a slight increase in the frequency of CD57⁺ γδ T cells during the first week, γδ T-cell composition remained stable, retaining a differentiated, senescent (CD57⁺) phenotype together with a highly cytotoxic profile (Figures 3C and S3C). A moderate increase in PD-1 expression was observed during the first week (15% to 20%), without concomitant upregulation of other inhibitory receptors such as CD39 or Tim-3. In contrast, CD69 expression was significantly increased at Day 4 in the majority of patients. This activation was transient, with CD69 expression returning toward baseline by Weeks 2–4 (Figures 3C and S3C).

**Figure 3.**
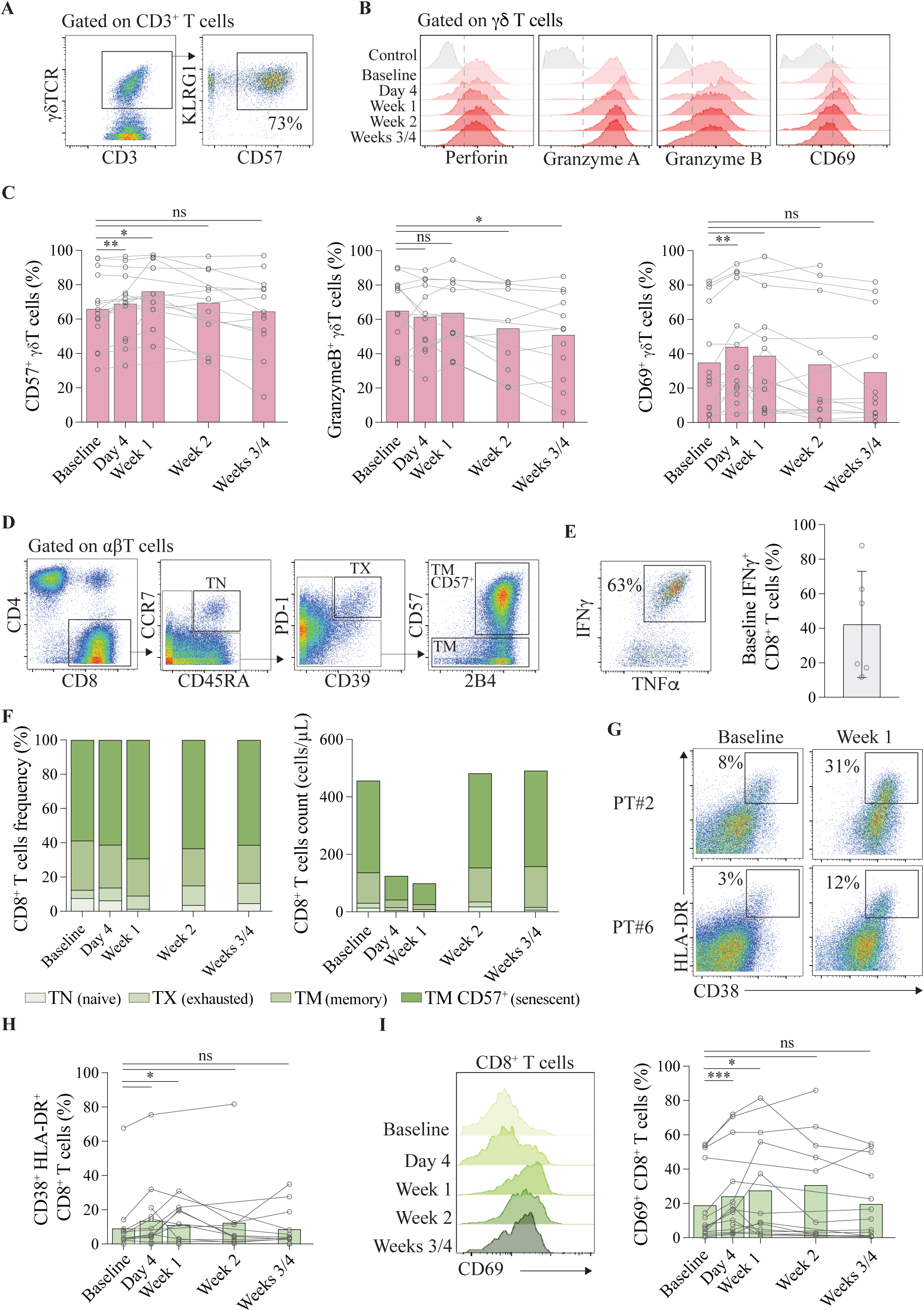
Activation of γδ and CD8^+^ T cells occur in the first week of CD3xBCMA BsAb therapy. **A.** Gating strategy used to characterize γδ T cells phenotype and their CD57 expression by mass cytometry. Representative data from one patient. **B.** Histograms of markers expression by γδ T cells during CD3xBCMA BsAB therapy. Representative data from one patient compared to its naive CD8^+^ T cells (Control). **C.** Frequencies of senescent (CD57^+^), cytotoxic (GrB^+^) and activated (CD69^+^) γδ T cells during CD3xBCMA BsAb therapy. **D.** Gating strategy used to identify CD8^+^ T cells main differentiation states (TN, TX, TM and TM CD57^+^). Representative data from one patient. **E.** Expression of IFNγ in CD8^+^ T cells after (left) and frequency of IFNγ^+^ CD8^+^ T cells from patients (right) at baseline. Data from mass cytometry after PMA/ionomycin stimulation. **F.** Frequency (left panel) and count (right panel) of CD8^+^ T cells main differentiation states in patients PBMCs during CD3xBCMA BsAb therapy (n=14). **G.** Gating strategy used to identify CD38^+^HLA-DR^+^ CD8^+^ T cells. **H.** Frequency of CD38^+^HLA-DR^+^ CD8^+^ T cells during CD3xBCMA BsAb therapy. **I.** Histograms of CD69 expression by CD8^+^ T cells during CD3xBCMA BsAB therapy. Representative data from one patient (left panel). Frequency of activated (CD69^+^) CD8^+^ T cells during CD3xBCMA BsAb therapy (right panel). Paired t-tests. ns= non-significant *p ≤ 0.05, **p ≤ 0.01, ***p ≤ 0.001.

According to the recently proposed guidelines for T-cell nomenclature^16^ and using a hierarchical gating strategy, the major conventional CD8⁺ T-cell differentiation states were defined as naïve (TN), exhausted (TX), memory (TM CD57⁻), and senescent memory (TM CD57⁺) (Figure 3D; see Methods). At baseline, CD8⁺ T cells exhibited preserved functionality, as demonstrated by their ability to produce IFNγ after *in vitro* stimulation together with high expression of cytotoxic molecules (Figures 3E and S4A). Approximately 25% of CD8⁺ T cells corresponded to TM CD57⁻ cells, which expressed high levels of KLRG1 and low levels of CD127, consistent with late effector memory T cells endowed with potent cytotoxic function (Figures 3F and S4B). TM CD57⁺ cells accounted for approximately 50% of CD8⁺ T cells and represented a highly differentiated, senescent memory population characterized by limited proliferative capacity while retaining strong cytotoxic function (Figures 3F and S4B). As observed for γδ T cells, absolute CD8⁺ T-cell counts decreased after treatment initiation across all differentiation subsets. Following recovery to baseline by Week 2, CD8⁺ T cells maintained a stable late effector/senescent differentiation profile (Figures 3F and S4C). Similarly, upregulation of inhibitory receptors, including PD-1, remained limited (<10%) and transient. By contrast, co-expression of CD38 and HLA-DR, a hallmark of recently activated T cells, increased significantly during the first week (from approximately 10% to 15% on average). Furthermore, CD69 expression increased significantly by 2-fold at Day 4 and remained elevated during the first 1–2 weeks, with marked inter-patient variability (6%–80%). These early activation-associated phenotypic changes were transient and returned to baseline by Weeks 2–4 (Figures 3G–I, S4D).

Within the CD4⁺ compartment at baseline, regulatory T cells (Tregs) represented approximately 10% of total CD4⁺ T cells and displayed high CD39 expression (60–90%), consistent with a suppressive phenotype (Figures 4A–C). Non-Treg CD4⁺ T cells were predominantly Th1- polarized (IFNγ⁺), with approximately 40% corresponding to TM CD57⁻ cells expressing high CD127 and low KLRG1, indicative of functional competence and long-term survival potential (Figures 4B, 4D, and S5A). CD4 TM CD57⁺ cells accounted for approximately 20% of CD4⁺ T cells and displayed cytotoxic characteristics without expression of NK-associated receptors (Figures 4B and S5B–D). As observed for γδ and CD8⁺ T cells, absolute CD4⁺ T-cell counts decreased following treatment initiation across all subsets and recovered by Week 2 (Figures 4B and S6A). Although the proportion of Tregs transiently increased at Day 4 to approximately 15%, the relative composition of CD4⁺ T-cell differentiation states remained stable (Figures 4B and S6A–B). Inhibitory receptor expression, including PD-1, remained low (<10%) and was only transiently increased. In contrast to γδ and CD8⁺ T cells, activation markers (CD69, CD38, and HLA-DR) remained consistently low (<10%) throughout the first month, with only occasional cases of transient activation (Figures 4E and S6C).

**Figure 4.**
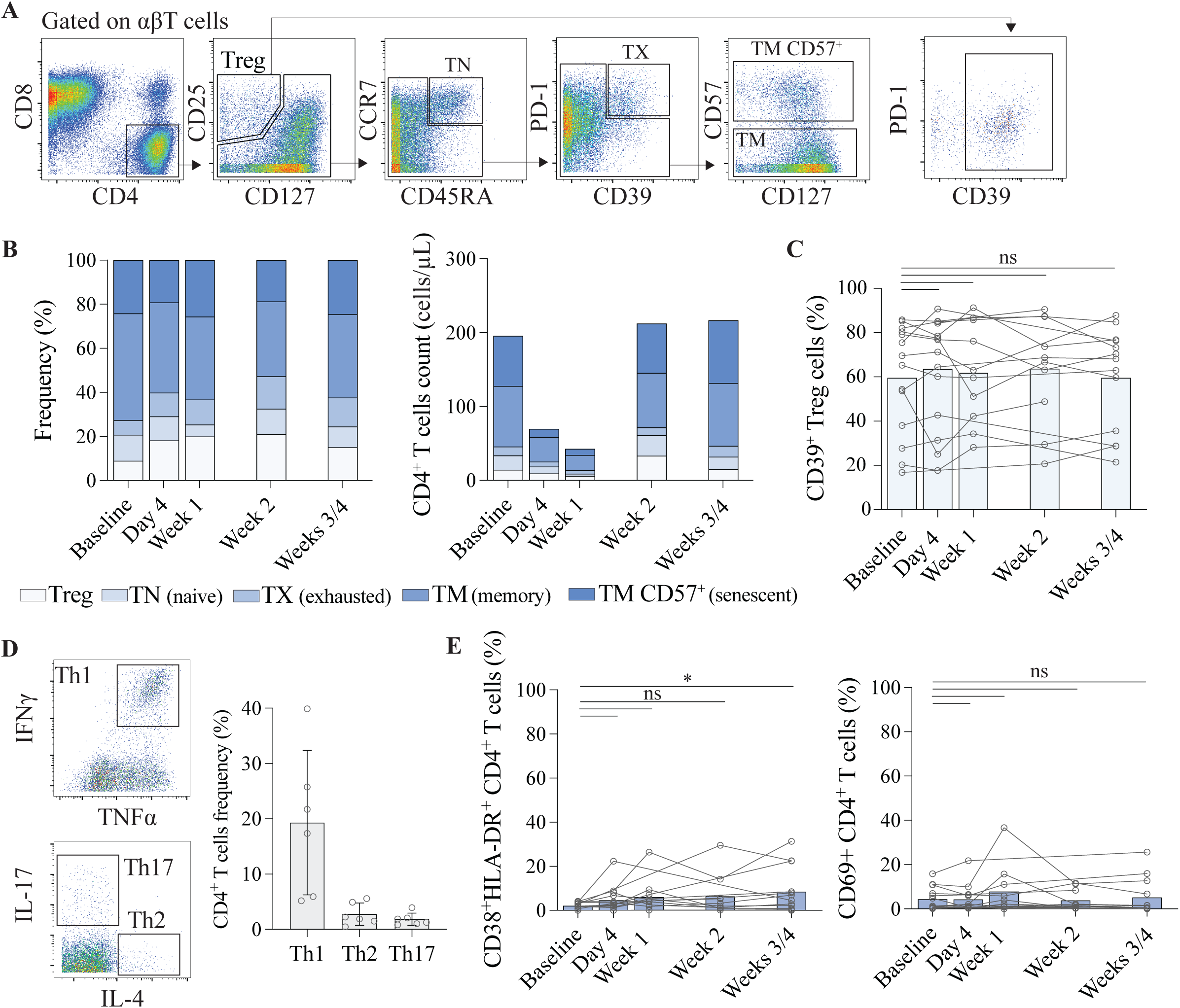
No CD4^+^ T cells activation occur during the first month of CD3xBCMA BsAb therapy. **A.** Gating strategy used to identify CD4^+^ T cells main differentiation states (Treg, TN, TX, TM and TM CD57^+^). Representative data from one patient. **B.**. Frequency (left panel) and absolute count (right panel) of CD4^+^ T cells main differentiation states in patients PBMCs during CD3xBCMA BsAb therapy. **C.** Frequency of CD39^+^ Treg cells during CD3xBCMA BsAb therapy. **D.** Polarization of CD4^+^ T cells from patients at baseline. Data from mass cytometry after pma/ionomycin stimulation. **E.** Frequencies of CD38^+^HLA-DR^+^ and CD69^+^ non-Treg CD4^+^ T cells during CD3xBCMA BsAb therapy. Paired t-tests. ns= non-significant *p ≤ 0.05, **p ≤ 0.01, ***p ≤ 0.001.

Collectively, these results show that during the first two weeks of treatment, subsets of γδ and CD8⁺ T cells undergo transient activation, characterized by increased expression of early activation markers (CD69, CD38/HLA-DR). Importantly, this heterogenous activation and kinetic between patients, occurs without major remodeling of T-cell differentiation states, cytotoxic potential, or plasmatic IFNγ/TNFα level. Notably, no comparable phenotypic changes were detected at later time points (Weeks 3–4), despite continued weekly administration of the bispecific antibody.

### Early lymphocyte dynamics, CXCL10 levels, sBCMA and bone lesions correlate with clinical response to treatment

To investigate the relationship between T-cell immune dynamics and clinical outcome, defined by immunochemical response after one month^17^, we integrated clinical and immunological variables using factorial analysis of mixed data (FAMD). FAMD enables the joint analysis of qualitative and quantitative variables within a single analytical framework. Our dataset comprised 14 clinical variables and 193 immunological features measured longitudinally in 14 patients throughout treatment (n = 5 timepoints) (Table S2). Because some variables contained missing values due to limited biological material, missing data were imputed prior to analysis. Qualitative variables were encoded as indicator matrices, as in multiple correspondence analysis (MCA), whereas quantitative variables were centered and scaled, as in principal component analysis (PCA). This approach allows variables with different scales and data types (i.e., percentages, cytokine concentrations, and clinical characteristics) to contribute equally to the analysis. FAMD results indicate that responders and non-responders formed distinct group, indicating that the integrated immune features captured differences associated with treatment response (Figure S7). A correlation network was subsequently generated by assessing the proximity of variables within this FAMD multidimensional space and retaining only the strongest positive and negative associations (Figure 5A and S8). Using this approach, we identified a cluster of immunological features associated with clinical response to treatment (Figure 5B).

**Figure 5.**
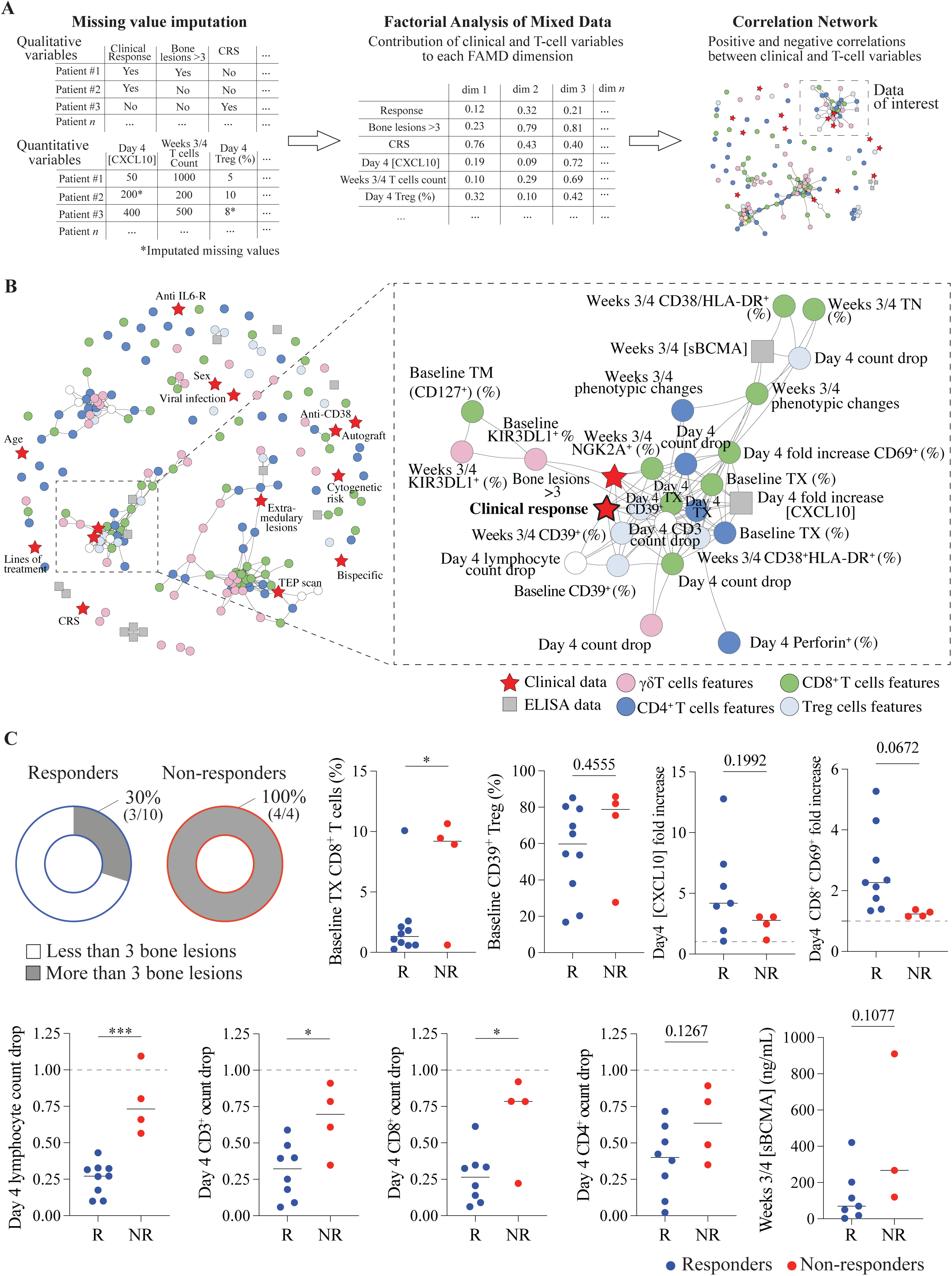
Integrating immunological and clinical data through Factorial Analysis of Mixed Data (FAMD). **A.** Schematic representation of FAMD and correlation network generation from imputation qualitative and quantitative data from patients. **B.** Correlation network of variables included in the FAMD analysis. n=14 patients. **C.** Verification of variables correlating with response to treatment in the correlation network without imputed values. Fold increase = Day 4/Baseline ratio. Count drop = Day 4/Baseline count ratio. Dotted line at y=1. Median, Fisher’s exact test (bone lesions), p =0.07/Unpaired t-tests. ns= non-significant *p ≤ 0.05, **p ≤ 0.01, ***p ≤ 0.001.

This cluster was predominantly composed of variables measured during the first week of therapy. To validate the contribution of each variable, we analyzed the raw data without imputed values. Some associations, particularly those involving KIR and NKG2A receptor expression, were no longer retained, most likely because of the reduced number of available observations (Figure S9). Compared with non-responders, responders tended to exhibit lower frequencies of exhausted T cells (TX) and CD39⁺ Tregs, together with fewer bone lesions at baseline (Figure 5C). Following treatment, responders were more likely to exhibit an increase in plasma CXCL10 levels, a more pronounced decrease in circulating lymphocyte counts, and a two-fold increase in CD69 expression at Day 4. Responders also exhibited more pronounced phenotypic remodeling of the T-cell compartment, as quantified by Jensen–Shannon divergence (JS div), and lower plasma sBCMA concentrations at later time points (Figures 5C and S9). Importantly, neither baseline nor treatment-induced T-cell differentiation states, including the frequency of senescent T cells, nor T-cell cytotoxic profiles were associated with clinical outcome. Likewise, no associations were observed with other clinical characteristics, including patient age, line of therapy, or the occurrence of cytokine release syndrome (CRS) (Figures 5B and S8).

To further validate these immunological and clinical correlates of treatment response, we analyzed an extended cohort comprising newly recruited baseline patients together with additional longitudinal plasma samples. In this extended cohort, a greater proportion of non-responders presented with more than three bone lesions (54.8% vs. 85.7% in non-responders) (Figure 6A). Likewise, only baseline frequencies of CD8⁺ TX cells remained significantly higher in non-responders (1.5% vs. 5.9%) (Figure 6B). We confirmed the higher sBCMA concentrations in non-responders, and longitudinal analyses demonstrated that this difference persisted throughout the first month of treatment (Figure 6C, S10A). Similarly, plasma CXCL10 concentrations were significantly lower in non-responders at Day 4 (336.0 pg/mL vs. 173.2 pg/mL), and longitudinal analyses confirmed persistently lower CXCL10 levels in non-responders during the first week of therapy (Figure 6D, S10B). Day 4 CXCL10 levels also significantly correlated with the early drop of lymphocytes (Figure S10C). Moreover, consistent with these observations, responders exhibited a significantly greater lymphocyte count drop at Day 4 and Week 1 compared to non-responders. As reported previously, responders also exhibited higher baseline circulating lymphocyte count (Figure 6E).

**Figure 6.**
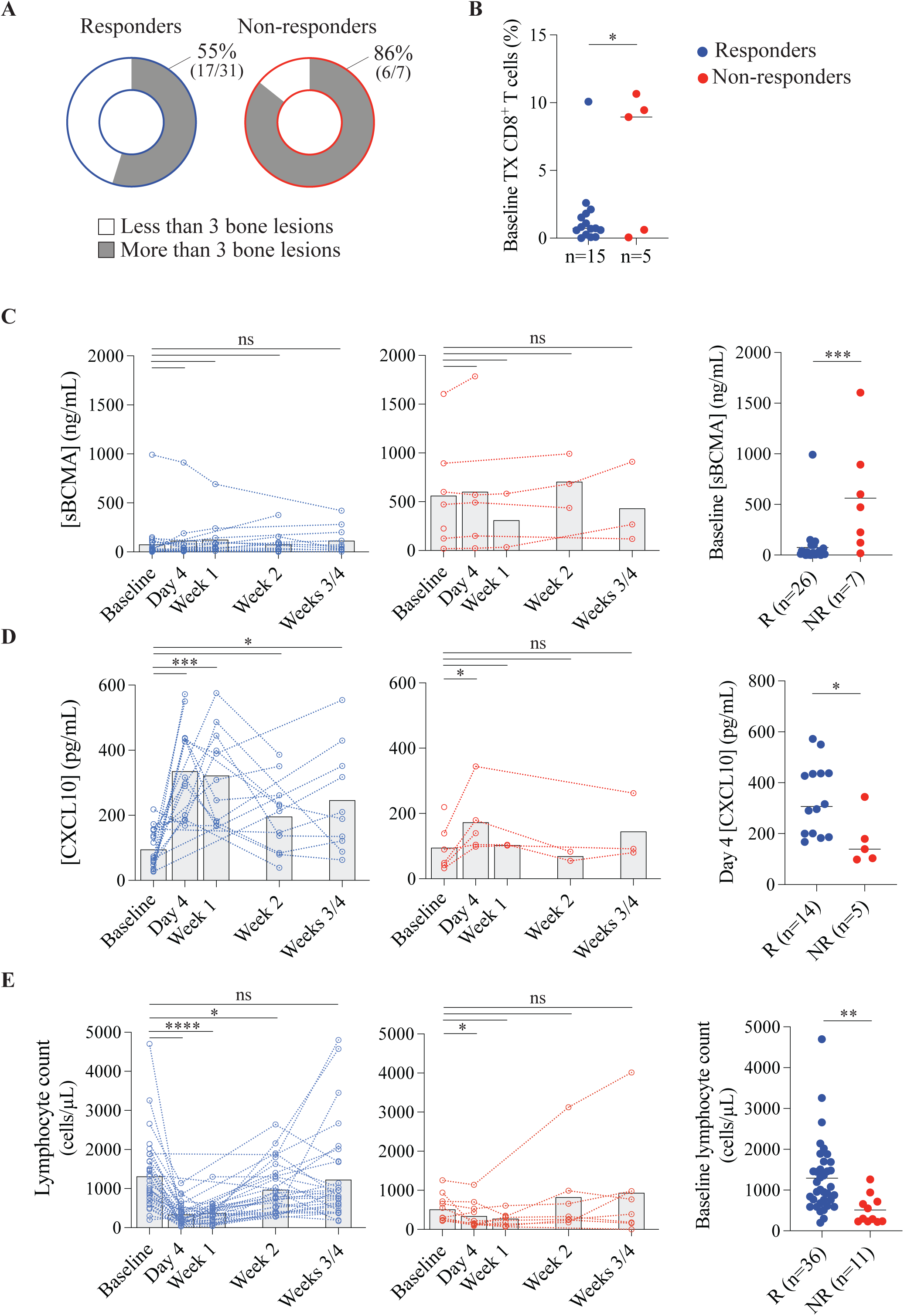
Confirmation of FAMD results and identification of potential biomarkers of response to CD3xBCMA BsAb therapy on an extended cohort. **A.** Pie chart of the distribution of patients with more than 3 bone lesions. Fisher’s exact test, p=0.21. **B.** Baseline TX CD8^+^ and CD4^+^ T cells frequency at baseline. **C.** Plasma concentrations of sBCMA during CD3xBCMA BsAb therapy (middle and left panels) and baseline sBCMA plasma concentrations (right panel). **D.** Plasma concentrations of CXCL10 during CD3xBCMA BsAb therapy (middle and left panels) and Day 4 CXCL10 plasma concentrations (right panel). **E.** Lymphocyte count during CD3xBCMA BsAb therapy (middle and left panels) and baseline lymphocyte count (right panel). (n=18 to n=47 patients). Unpaired t-tests/Paired t-tests, ns= nonsignificant *p ≤ 0.05, **p ≤ 0.01, ***p ≤ 0.001 ****p ≤ 0.0001. R= Responders; NR=Nonresponders.

Together, these findings demonstrate that clinical response to CD3×BCMA therapy is associated with an early immune response occurring within the first week of treatment rather than with major changes in T-cell differentiation. Number of bone lesions at baseline, early CXCL10 induction, profound but transient lymphocyte count drop, and low baseline sBCMA levels emerge as robust correlates of therapeutic efficacy and represent promising biomarkers for early treatment monitoring.

## Discussion

Beyond the strong intra and inter-patients’ heterogeneity measured, our observations support a biological model in which initial exposure to CD3×BCMA bispecific antibody (BsAb) triggers an acute priming phase driven by CD3-mediated crosslinking of T cells with BCMA-expressing tumor cells in the bone marrow. This interaction induces rapid T-cell activation, leading to the release of inflammatory chemokines CXCL10. As a key ligand of CXCR3, CXCL10 promotes the recruitment of activated T cells to inflamed tissues. Consistent with this mechanism, preclinical studies using a CD20×CD3 BsAb in a humanized mouse model have shown that early lymphopenia is associated with T-cell redistribution to tissues and increased CXCL10 levels^18^. In our cohort, the transient activation of γδ and CD8⁺ T cells observed between Day 4 and Week 2 supports a model in which CD3×BCMA BsAb therapy induces a short systemic immune response, characterized by increased expression of CD69, CD38, and HLA-DR, followed by restoration within the first month. The early decrease in lymphocyte count was already detectable 24 hours after treatment initiation, whereas our first immunophenotyping timepoint was Day 4. This observation suggests that the immune activation observed at Day 4 may represent an established phase of a response that is initiated even earlier, potentially within the first 24 hours following treatment initiation. The transient activation may reflect the recirculation of activated T cells between the bone marrow and blood during the early phase of tumor cell elimination. The recovery of lymphocyte counts by Week 1, together with the transient nature of T-cell activation despite continued weekly dosing, suggests that this response is largely confined to the first-dose pharmacodynamic effect, after which T-cell distribution and activation reach a new equilibrium. The absence of recurrent lymphocyte drops or sustained T-cell activation with subsequent administrations indicates that later doses primarily maintain target engagement rather than re-induce strong systemic immune activation. These findings indicate that CD3×BCMA BsAb therapy induces a rapid, first-dose–dominated immune activation phase within the first week, with important implications for dosing strategies. This supports the rationale for response-adapted dosing approaches after the first week, hypothesizing that sustained clinical efficacy still depends on maintaining adequate drug exposure over time^19^.

Although integrated data analysis using FAMD enabled a comprehensive two-dimensional visualization of relationships within this complex dataset, handling missing data remains a critical limitation for biological interpretation. Imputation can introduce artificial structure that does not necessarily reflect true biological signals. Therefore, the interpretation of the derived correlation network must be approached with caution, ensuring that observed patterns are biologically plausible rather than mathematical artifacts. In our study, variables with more than 30% missing values were excluded to preserve interpretability^20^. Maintaining a low proportion of missing data is essential to preserve the robustness and biological relevance of the resulting representation. Beyond its biological findings, our study illustrates the potential of FAMD as an integrative method to jointly analyze heterogenous clinical and immunological variables to provides a coherent data network visualization. Our study identifies a combination of baseline clinical characteristics and early treatment-induced immune changes that consistently associate with therapeutic efficacy. Specifically, a lower number of bone lesions at baseline, low baseline sBCMA levels, rapid induction of plasma CXCL10, and a profound but transient decline in circulating lymphocyte counts during the first week of treatment emerged as coordinated features of responding patients. While patients with extensive disease burden, including more high number of bone lesions, have previously been reported to experience inferior outcomes following BCMA- directed T-cell therapies^21^, and decline in circulating lymphocyte counts during the first week^22^, our findings suggest that integrating bone lesions with dynamic immune biomarkers may substantially improve early prediction of treatment response. Overall, these findings suggest that clinical response to CD3xBCMA BsAb therapy is driven by a pre-existing state of immune fitness characterized by a larger and more functional T cell compartment. T cells in this context appear less exhausted and more capable of mounting a rapid and dynamic response at the tumor site shortly after the first dose. In contrast with previous reports^9,13^, we did not observe significant differences between responders and non-responders in several immune subsets that have been associated with clinical outcome, including CD38^+^ Treg and effector CD8^+^ T cells frequencies. This discrepancy may reflect differences in patient populations, treatment history, or analytic approaches across studies. The limited size of our cohort also reduces the statistical power to detect more subtle immunological differences. We analyzed several time points for each patient in both plasma and PBMC, so our study provides a very deep analysis of each patient, rather than a shallow analysis of many patients^23^. Future efforts should focus on confirming and combining biomarkers of response into a multiparametric immunoscore capable of capturing the complexity and heterogeneity of patient’s immune landscape. Beyond their relevance for current CD3xBCMA BsAb therapies, it will be important to determine whether the early immune dynamics identified here are shared across other T cell engagers therapies approved for cancers and autoimmune diseases^24^. Finally, these findings may also have implications for the development and clinical implementation of next-generation trispecific antibodies, where the pre-existing immune fitness is likely to remain a key determinant of treatment outcome^25^.

## Material and methods

### Study design

Blood aspirates were collected from refractory/relapsed multiple myeloma patients before and after their first injection of CD3xBCMA BsAb therapy (teclistamab or elranatamab). In the dose-escalation step, patients received three injections of increasing concentration of CD3xBCMA BsAb in the first week. Patients were then treated weekly CD3xBCMA BsAb. The use of human samples was approved by the appropriate institutional review boards and ethical committees (CPP 2015-08-11DC).

### Cell and plasma isolation

Blood samples were centrifuged for 10min at 400g to collect plasma and store it at −70°C. PBS was added to the remaining blood sample in a volume to match the volume of plasma retrieved. The sample was then carefully layered on a Ficoll density gradient and centrifuged at 400g for 15min without brake. PBMC ring was collected from the interface and washed with PBS at 400g for 5min. Cells were cryopreserved in anonymized cryotubes using 90% fetal bovine serum (FBS) and 10% DMSO and stored in liquid nitrogen.

### Mass cytometry sample processing

Antibodies were conjugated to their given metal isotopes in-house according to the manufacturer’s protocol (Standard BioTools). Cryopreserved samples were thawed at 37°C and washed in PBS. Samples were prepared and stained as previously described^26,27,28,29^. In brief, cells were stained for 10min at room temperature (RT) with a surface antibody cocktail in staining buffer (PBS + 0.5% BSA + 0.02% sodium azide+ 250 μg/mL DNAse) followed by 5μM cisplatin for 5min for viability staining. Cells were then permeabilized for 30min at RT using Cyto-Fast Fix/Perm solution (BioLegend), washed and stained with an intracellular antibody cocktail in permeabilization buffer for 30min at RT. After final washes in permeabilization and staining buffers, cells were fixed in 2% paraformaldehyde (PFA) overnight. On the day of CyTOF acquisition, cells were washed in H_2_O and DNA staining was performed using 250nM Cell-ID Intercalator-Iridium (Standard BioTools) in H_2_O for 5min (see supplementary table 3 for antibody clones and metal conjugates).

### PMA/ionomycin stimulation

For cytokine detection, samples were pre-stained with anti-CD69, then stimulated for 4 hours at 37°C with PMA (50 ng/mL), ionomycin (1 μg/mL), and brefeldin A (1 μg/mL) in RPMI + 10% FBS. Cells were then stained for surface and intracellular markers as previously mentioned. (see supplementary table 3 for antibody clones and metal conjugates).

### Mass cytometry data analysis

On CyTOF data (.fcs file), using the R flowCore package (v2.12.2) in RStudio, zero values were replaced with a random uniform distribution between 0 and −1 to improve data visualization in dot plot representations. CyTOF data were analyzed using FlowJo software (v10.4). Among live immune cells (DNA⁺ cisplatin⁻ CD45⁺), monocytes (CD14⁺) and B cells (CD19⁺) were excluded by manual gating. According to the guidelines for T-cell nomenclature (v1.0) and using a hierarchical gating strategy, the main T-cell lineages (CD3⁺) were defined as follows: γδ T cells (γδTCR⁺), MAIT cells (TCR Vα7.2⁺ CD161⁺), and CD4⁺ or CD8⁺ T cells. CD8⁺ T-cell differentiation states were defined as follows: naïve (CD8 TN: CCR7⁺ CD45RA⁺), exhausted (CD8 TX: PD-1⁺ CD39⁺), effector (CD8 TM: CD57⁻ CD45RO⁺), and senescent (CD8 TM CD57⁺). Similarly, within CD4⁺ T cells, differentiation states were defined using a hierarchical gating strategy as follows: regulatory T cells (Treg: CD25⁺ CD127low/−), naïve (CD4 TN: CCR7⁺ CD45RA⁺), exhausted (CD4 TX: PD-1⁺ CD39⁺), effector (CD4 TDM: CD57⁻ CD45RO⁺), and senescent (CD4 TDM CD57⁺). Following ex vivo stimulation, T helper cell functional polarization was defined as: TH1 (IFNγ⁺), TH2 (IL-4⁺), and TH17 (IL-17⁺).

UMAP analysis was performed on selected markers (Supplementary table 3) using the umap package (v0.2.10) on transformed data, with each patient analyzed independently or grouped when staining was performed in the same batch. The Jensen–Shannon divergence (JS div) was calculated as previously described^30^.

### Measurement of cytokine levels

CXCL12, CXCL10, Granzyme A, BCMA, TNFα, IFNγ and IL-6 levels were measured with DuoSet ELISA kits from Bio-Techne using the manufacturer protocol (DY350-05, DY266-05, DY2905-05, DY193, DY210-05, DY285B-05, DY206-05, respectively). Plates were read at 450nm and 570nm wave lengths using CLARIOstar. Data analysis was performed using GraphPad Prism (v10.1.1) and Sigmoidal 4PL standard curve interpolation.

### Factorial analysis of mixed data

Factorial analysis of mixed data was performed using the FactoMineR package (v2.11) in RStudio^31^. Since the algorithm cannot handle missing values, missing data were imputed using missMDA (v1.19)^20^, provided that the proportion of missing values for a given variable did not exceed 30%. A correlation network based on the first 10 FAMD dimensions was generated using the igraph package (v2.0.3), only displaying correlations with a threshold higher than 90%.

### Statistical analysis

CyTOF data was analyzed using FlowJo software (version 10.4). Statistical analyses were performed using GraphPad Prism (version 10, GraphPad Software). Comparisons between two groups were performed using either paired or unpaired t-tests. Comparisons between each timepoint to baseline were performed with mixed effect analysis tests and Dunnett’s multiple comparison test.

## Data availability

Data are available on flow-repository website: https://flowrepository.org/XXXX

## Authorship Contributions

N.D. performed the experiments, analyzed the data and wrote the paper. L.A., A.G., M.M, C.C., helped perform the experiments and reviewed the paper. N.D., and Y.S. developed R scripts for data analysis. I.B., J.D., O.K., L.W., M.F., P.F., R.B., N.C., D.B., provided samples and discussed data and reviewed the paper. D.S-B., A.A., C.A. discussed the data and gave insightful feedbacks. M.V. and Y.S. Led, initiated and designed the project, performed the experiments, analyzed the data and wrote the paper.

---

**Supplementary figure 1.**
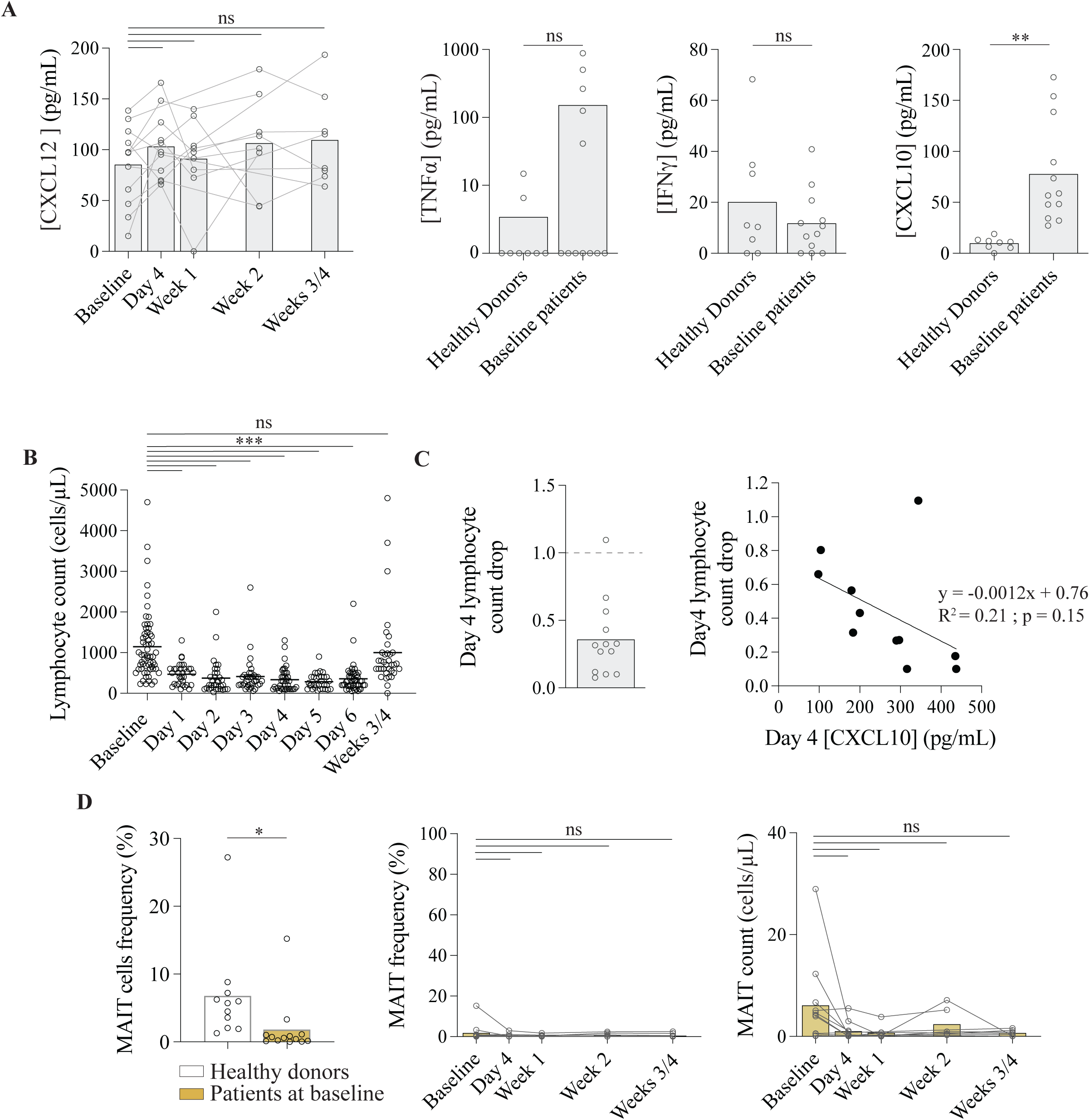
**A.** Plasma concentrations of CXCL12 during CD3xBCMA BsAb therapy (left panel) and plasma concentrations of TNFα, IFNγ and CXCL10 of healthy donors and patients at baseline (right panels). **B.** Lymphocyte count in the first 6 days and at Weeks 3/4 of treatment. n=33 to 50. **C.** Correlation between Day 4 lymphocyte count drop (Day 4/Baseline count ratio) and Day 4 CXCL10 concentration. Simple linear regression. **D.** MAIT cells frequency in healthy donors and patients at baseline (left panel). Frequency (middle panel) and count (right panel) of MAIT cells during CD3xBCMA BsAb therapy. Unpaired t-tests/Paired t-tests, ns= non-significant *p ≤ 0.05, **p ≤ 0.01, ***p ≤ 0.001.

**Supplementary figure 2.**
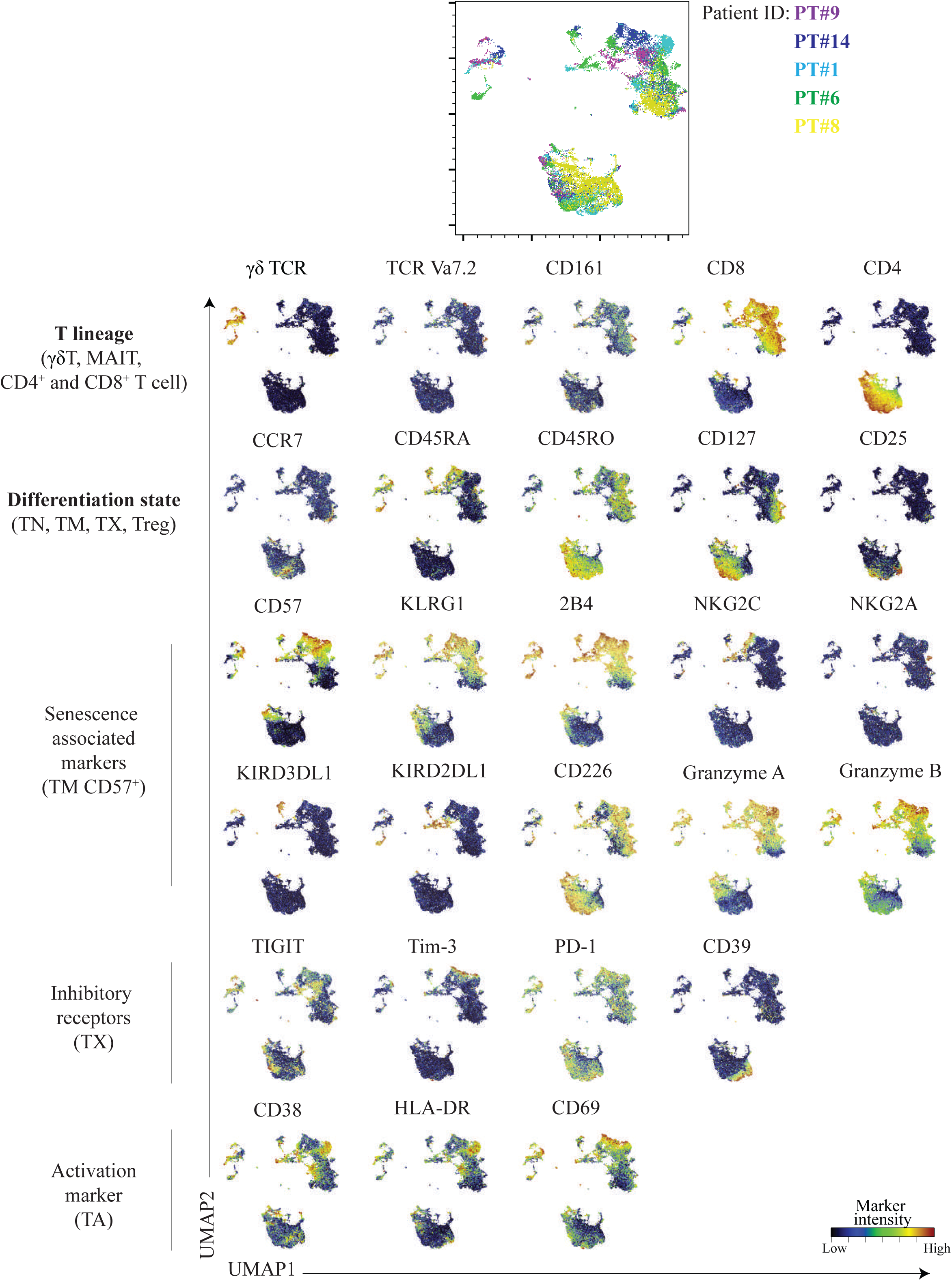
UMAP analysis of CD3⁺ T cells from patients PBMCs before CD3xBCMA BsAb therapy. Combined view (upper panel). Normalized marker expression intensities were calculated and overlaid on the UMAP plot (lower panels).

**Supplementary figure 3.**
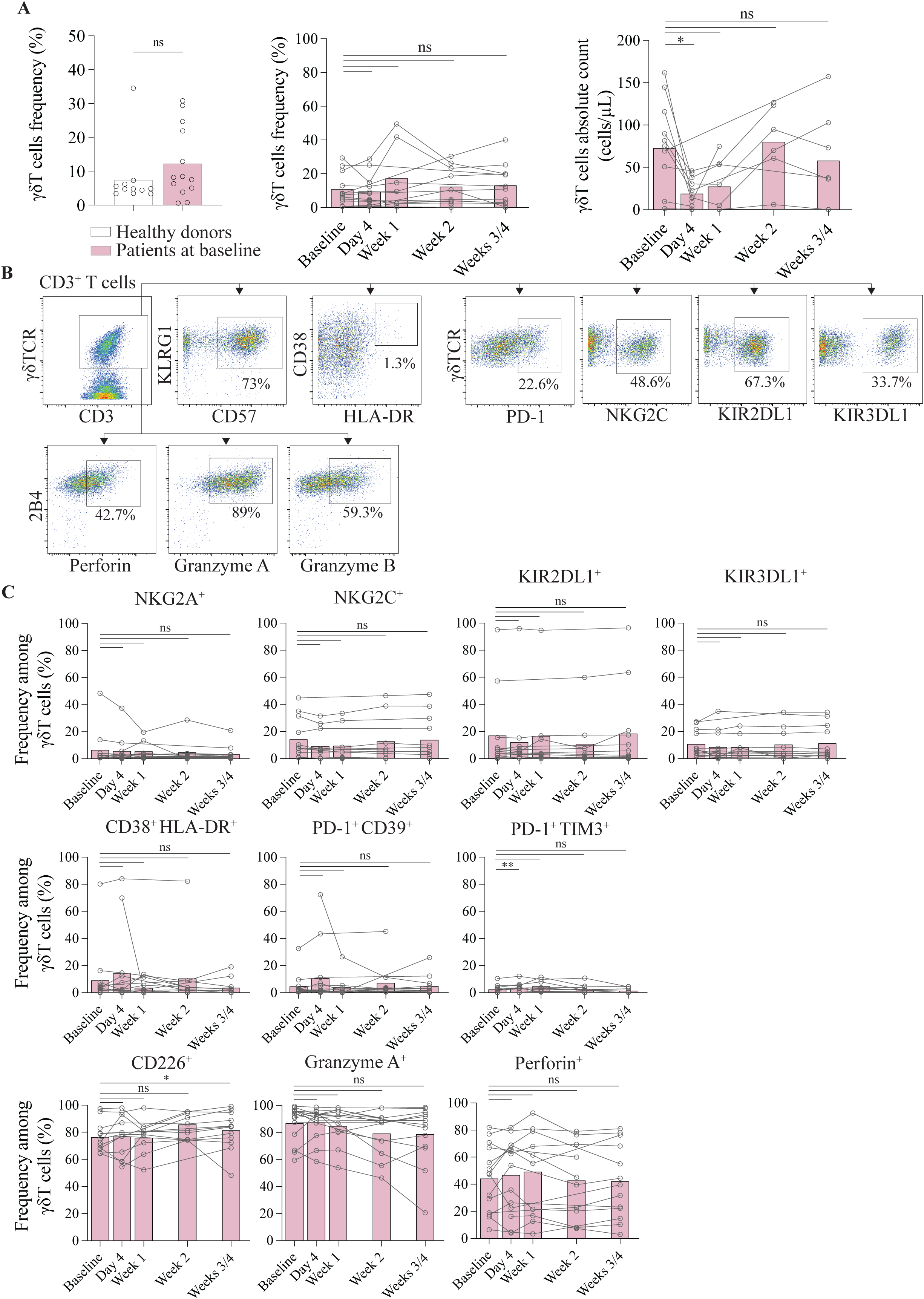
**A.** γδ T cells frequency in healthy donors and patients at baseline (left panel). Frequency (middle panel) and count (right panel) of γδ T cells during CD3xBCMA BsAb therapy. **B.** Gating strategy used to characterize γδ T cells phenotype. Representative data from one patient. **C.** Frequency of γδ T cells positive for various markers during CD3xBCMA BsAb therapy. Unpaired t-test/Paired t-tests, ns= non-significant *p ≤ 0.05, **p ≤ 0.01, ***p ≤ 0.001.

**Supplementary figure 4.**
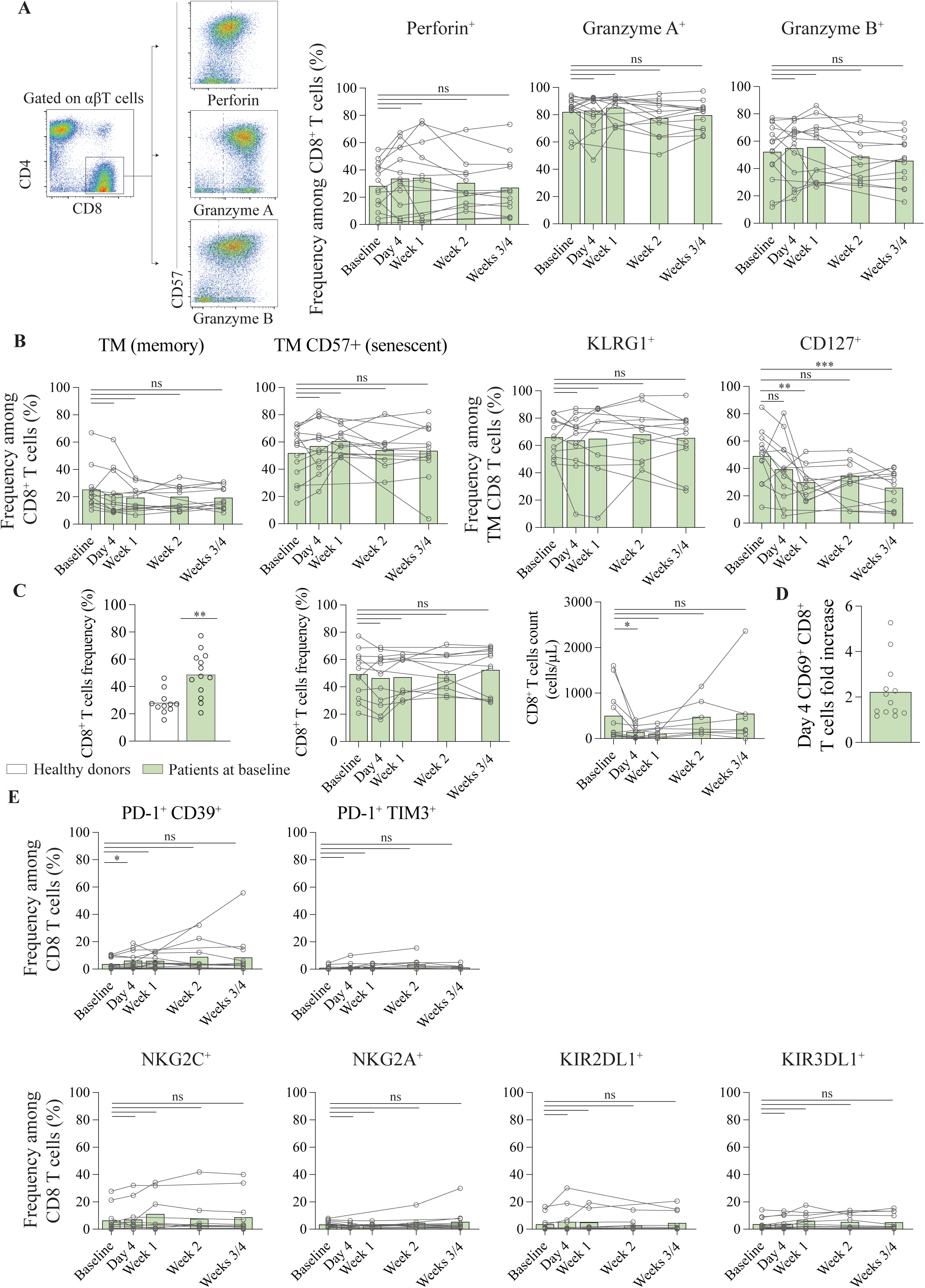
**A.** Gating strategy used to characterize CD8^+^ T cells cytotoxic profiles (left panel). Representative data from one patient. Frequency of Perforin, Granzyme A and Granzyme B CD8^+^ T cells during CD3xBCMA BsAb therapy (right panels). **B.** Frequency of TM and TM CD57^+^ CD8^+^ T cells during CD3xBCMA BsAb therapy (left panels). Frequencies of KLRG1^+^ and CD127^+^ in TM CD8^+^ T cells during CD3xBCMA BsAb therapy (right panels). **C.** CD8^+^ T cells frequency in healthy donors and patients at baseline (left panel). Frequency (middle panel) and count (right panel) of CD8^+^ T cells during CD3xBCMA BsAb therapy. **D.** Day 4 CD69^+^ CD8^+^ T cells fold increase (Day 4/Baseline ratio). **E.** Frequency of CD8^+^ T cells positive for various markers during CD3xBCMA BsAb therapy. Unpaired t-test/Paired t-tests, ns= non- significant *p ≤ 0.05, **p ≤ 0.01, ***p ≤ 0.001.

**Supplementary figure 5.**
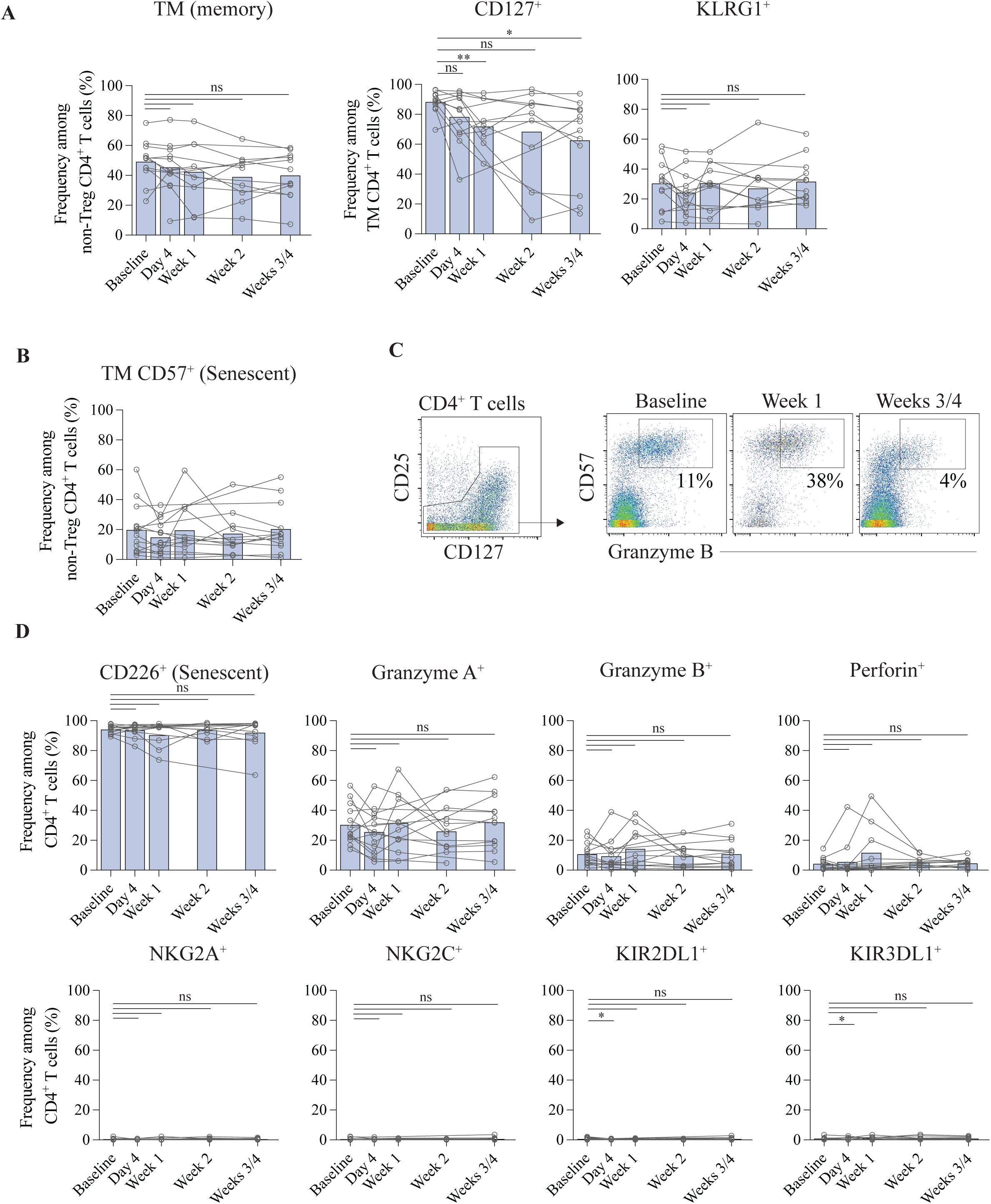
**A.** Frequency of TM CD4^+^ T cells (left panel) and frequency of CD127^+^ and KLRG1^+^ in TM CD4^+^ T cells during CD3xBsAb therapy (right panels) **B.** Frequency of TM CD57^+^ CD4^+^ T cells during CD3xBsAb therapy for various markers during CD3xBCMA BsAb therapy. **C.** Gating strategy used to identify Granzyme B^+^ CD4^+^ T cells. Representative data from one patient. **D.** Frequency of CD4^+^ T cells positive for various markers during CD3xBCMA BsAb therapy. Paired t-tests, ns= non-significant *p ≤ 0.05, **p ≤ 0.01, ***p ≤ 0.001.

**Supplementary figure 6.**
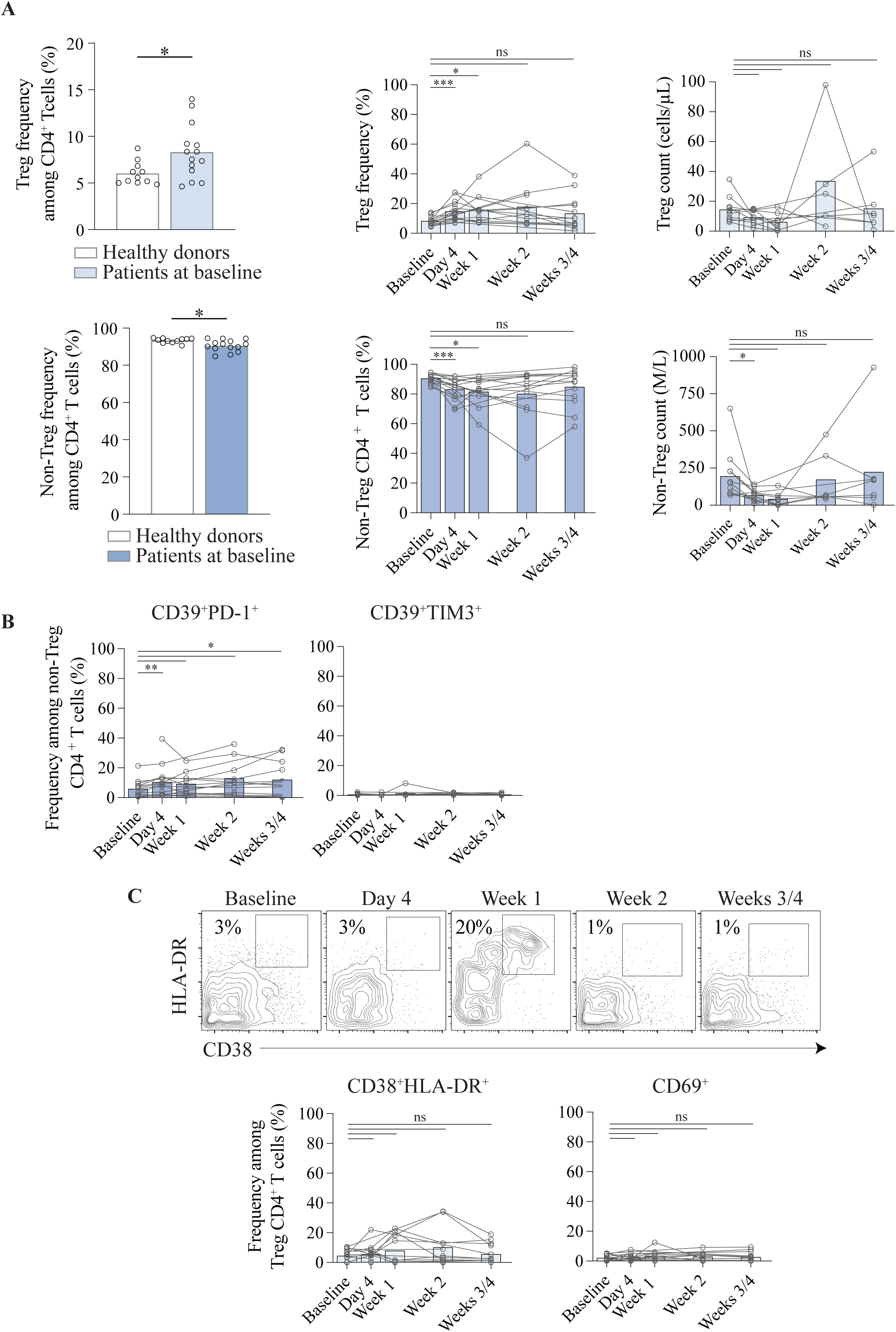
**A.** Treg cells frequency in healthy donors and patients at baseline (upper left panel). Frequency (upper middle panel) and count (upper right panel) of Treg T cells during CD3xBCMA BsAb therapy. Non-Treg cells frequency in healthy donors and patients at baseline (lower left panel). Frequency (lower middle panel) and count (lower right panel) of non-Treg cells during CD3xBCMA BsAb therapy. **B.** Frequency of CD4^+^ T cells positive for various inhibitory markers during CD3xBCMA BsAb therapy. **C.** Gating strategy used to identify Treg cells positive for activation markers (CD38 and HLA-DR). Representative data from one patient. (upper panel). Frequency of CD38^+^HLA-DR^+^ and CD69^+^ Treg cells during CD3xBCMA BsAb therapy. Unpaired t-test/Paired t-tests, ns= non-significant *p ≤ 0.05, **p ≤ 0.01, ***p ≤ 0.001.

**Supplementary figure 7.**
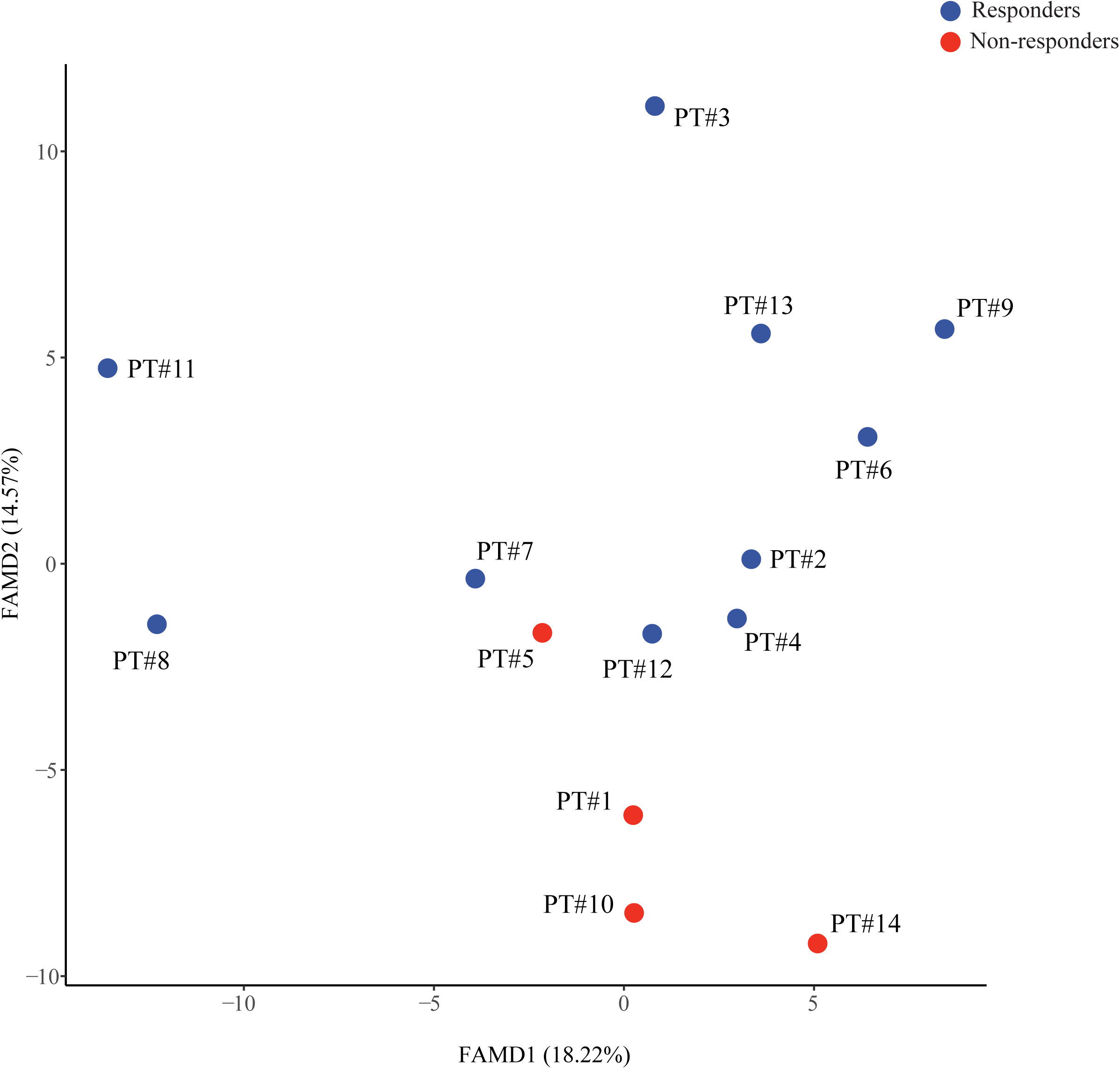
FAMD projection of responders and non-responders. n=14.

**Supplementary figure 8.**
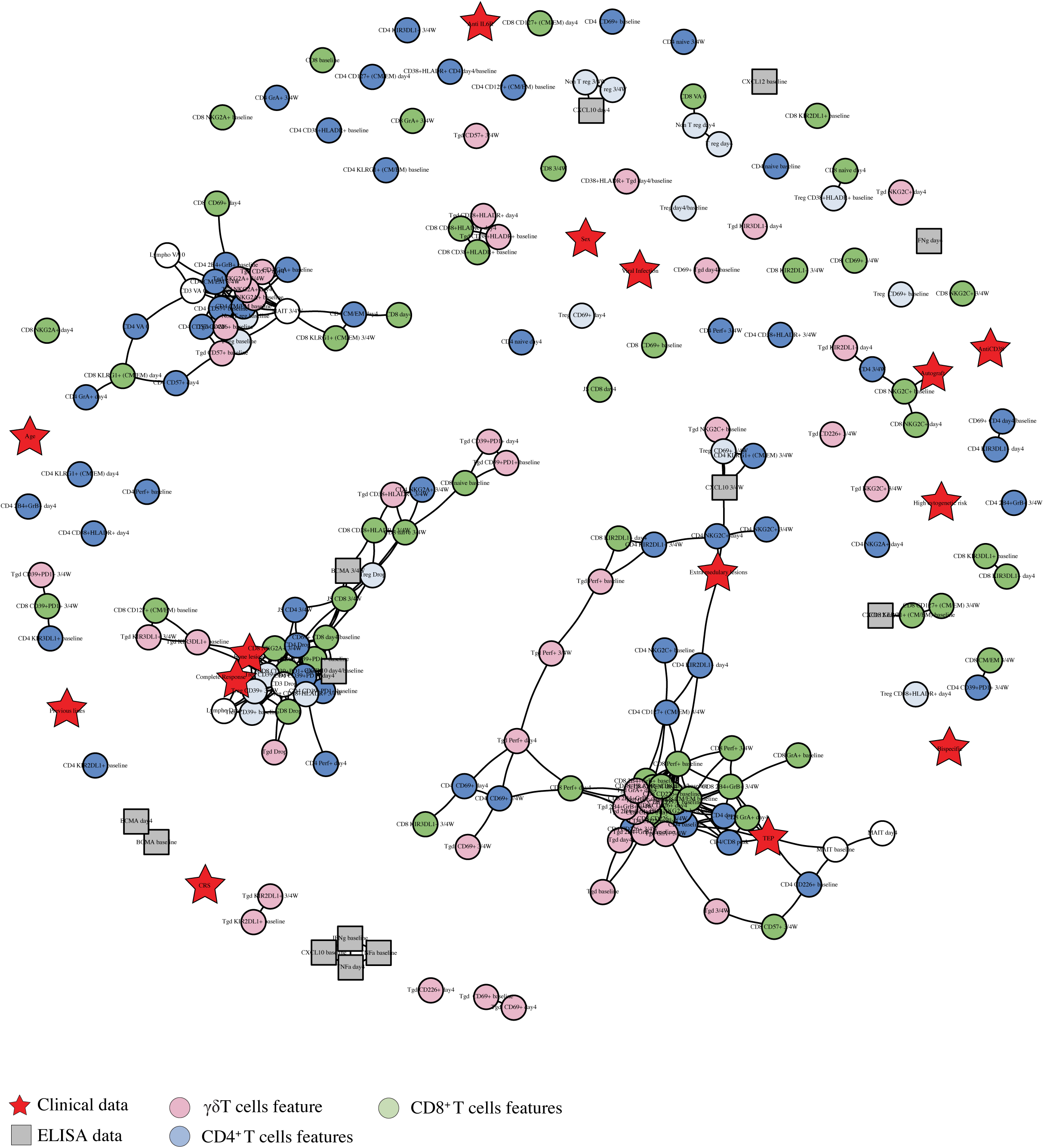
Complete correlation network of variables included in the FAMD analysis.

**Supplementary figure 9.**
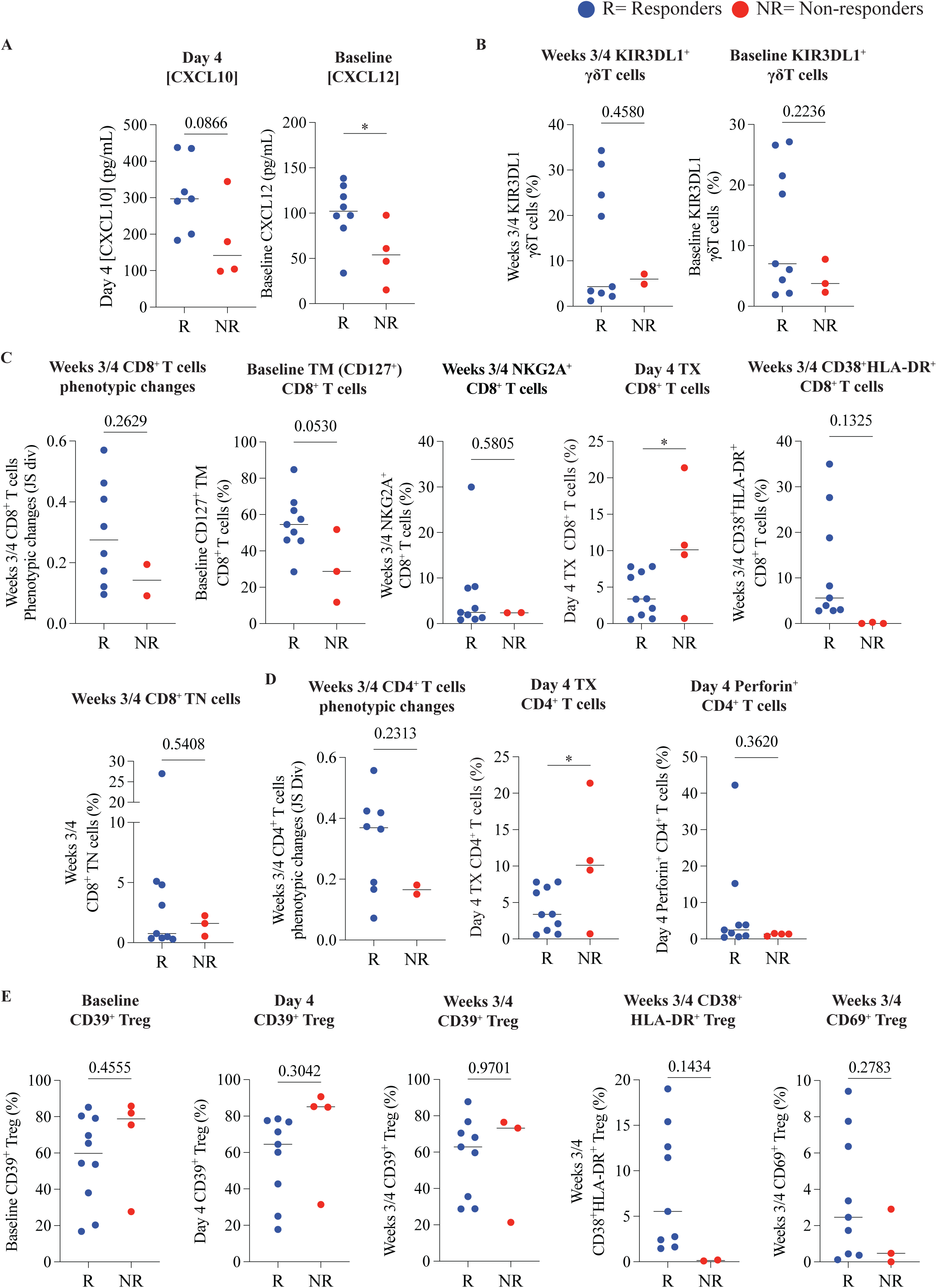
Verification of variables correlating with response to treatment in the correlation network without imputed values. **A.** Plasma cytokines-related features. **B.** γδ T cells-related features. **C.** CD8^+^ T cells-related features. **D.** CD4^+^ T cells-related features. **E.** Treg-related features. Median, Fisher’s exact test/Unpaired t-tests. ns= non-significant *p ≤ 0.05, **p ≤ 0.01, ***p ≤ 0.001.

**Supplementary figure 10.**
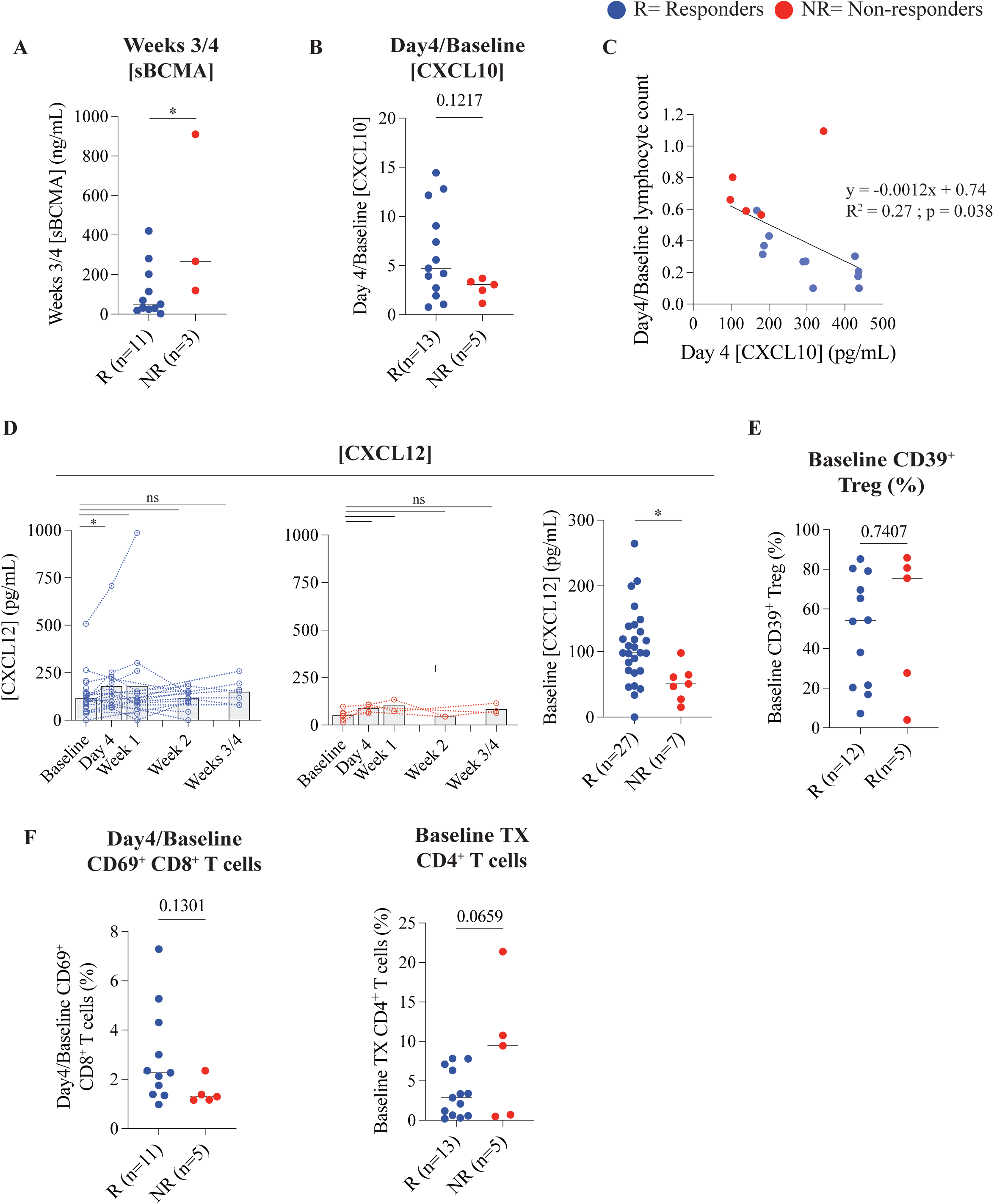
**A.** Weeks 3/4 sBCMA plasma concentration. **B.** Day4/Baseline CXCL10 plasma concentration. **C.** Correlation between Day 4 CXCL10 plasma concentration and Day 4/Baseline lymphocyte count. Simple linear regression. **D.** CXCL12 plasma concentration during CD3xBCMA BsAb therapy. **E.** Baseline CD39^+^ Treg frequency. **F.** Day 4/Baseline CD69^+^ CD8^+^ T cells frequency (left panel). Baseline TX CD4^+^ T cells frequency (right panel). Unpaired t-tests. ns= non-significant *p ≤ 0.05, **p ≤ 0.01, ***p ≤ 0.001.

## References

1. Shah, U. A. & Mailankody, S. Emerging immunotherapies in multiple myeloma. BMJ m3176 (2020) doi:10.1136/bmj.m3176.

2. Rajkumar, S. V. et al. International Myeloma Working Group updated criteria for the diagnosis of multiple myeloma. Lancet Oncol. 15, e538–e548 (2014).

3. Cowan, A. J. et al. Diagnosis and Management of Multiple Myeloma: A Review. JAMA 327, 464 (2022).

4. Hemminki, K., Hemminki, J., Försti, A. & Sud, A. Survival in hematological malignancies in the Nordic countries through a half century with correlation to treatment. Leukemia 37, 854–863 (2023).

5. Raab, M. S. et al. Multiple myeloma: practice patterns across Europe. Br. J. Haematol. 175, 66–76 (2016).

6. Moreau, P. et al. Teclistamab in Relapsed or Refractory Multiple Myeloma. N. Engl. J. Med. 387, 495–505 (2022).

7. Bahlis, N. J. et al. Elranatamab in relapsed or refractory multiple myeloma: the MagnetisMM- 1 phase 1 trial. Nat. Med. 29, 2570–2576 (2023).

8. Lee, H. et al. Impact of soluble BCMA and non–T-cell factors on refractoriness to BCMA- targeting T-cell engagers in multiple myeloma. Blood 144, 2637–2651 (2024).

9. Cortes-Selva, D. et al. Correlation of immune fitness with response to teclistamab in relapsed/refractory multiple myeloma in the MajesTEC-1 study. Blood 144, 615–628 (2024).

10. Lee, H. et al. Mechanisms of antigen escape from BCMA- or GPRC5D-targeted immunotherapies in multiple myeloma. Nat. Med. 29, 2295–2306 (2023).

11. Truger, M. S. et al. Single- and double-hit events in genes encoding immune targets before and after T cell–engaging antibody therapy in MM. Blood Adv. 5, 3794–3798 (2021).

12. Friedrich, M. J. et al. The pre-existing T cell landscape determines the response to bispecific T cell engagers in multiple myeloma patients. Cancer Cell 41, 711–725.e6 (2023).

13. Firestone, R. S. et al. CD8 effector T cells enhance teclistamab response in BCMA- exposed and -naïve multiple myeloma. Blood Adv. 8, 1600–1611 (2024).

14. Verkleij, C. P. M. et al. T-Cell Characteristics Impact Response and Resistance to T- Cell–Redirecting Bispecific Antibodies in Multiple Myeloma. Clin. Cancer Res. 30, 3006–3022 (2024).

15. Hipp, S. et al. A novel BCMA/CD3 bispecific T-cell engager for the treatment of multiple myeloma induces selective lysis in vitro and in vivo. Leukemia 31, 1743–1751 (2017).

16. Masopust, D. et al. Guidelines for T cell nomenclature. Nat. Rev. Immunol. 26, 298–313 (2026).

17. Lin, Y. et al. Consensus guidelines and recommendations for the management and response assessment of chimeric antigen receptor T-cell therapy in clinical practice for relapsed and refractory multiple myeloma: a report from the International Myeloma Working Group Immunotherapy Committee. Lancet Oncol. 25, e374–e387 (2024).

18. Cremasco, F. et al. Cross-linking of T cell to B cell lymphoma by the T cell bispecific antibody CD20-TCB induces IFNγ/CXCL10-dependent peripheral T cell recruitment in humanized murine model. PLOS ONE 16, e0241091 (2021).

19. Guo, Y. et al. Teclistamab Dosing in Responders: Modeling and Simulation Results from the MajesTEC-1 Study in Relapsed/Refractory Multiple Myeloma. Target. Oncol. 20, 651–661 (2025).

20. Josse, J. & Husson, F. **missMDA** : A Package for Handling Missing Values in Multivariate Data Analysis. J. Stat. Softw. 70, (2016).

21. Steinhardt, M. J. et al. Activity of CAR-T cells and bispecific antibodies in multiple myeloma with extramedullary involvement. Blood Cancer J. 15, 126 (2025).

22. Dettmar, K. et al. Transient lymphocyte decrease due to adhesion and migration following catumaxomab (anti-EpCAM x anti-CD3) treatment in vivo. Clin. Transl. Oncol. 14, 376–381 (2012).

23. Shasha, C. et al. Hallmarks of T-cell exhaustion and antigen experience are absent in multiple myeloma from diagnosis to maintenance therapy. Blood 145, 3113–3123 (2025).

24. Albayrak, G., Wan, P. K.-T., Fisher, K. & Seymour, L. W. T cell engagers: expanding horizons in oncology and beyond. Br. J. Cancer 133, 1241–1249 (2025).

25. Li, J. et al. YMN-V115: a novel humanized BCMA/GPRC5D/CD3 trispecific antibody in relapsed/refractory multiple myeloma. J. Immunother. Cancer 14, e013986 (2026).

26. Simoni, Y. et al. Bystander CD8+ T cells are abundant and phenotypically distinct in human tumour infiltrates. Nature 557, 575–579 (2018).

27. Li, S. et al. Bystander CD4^+^ T cells infiltrate human tumors and are phenotypically distinct. OncoImmunology 11, 2012961 (2022).

28. Aziez, L. et al. Pro-inflammatory role of granzyme K producing bystander CD8^+^ T cells in acute myeloid leukemia. Preprint at 10.1101/2025.08.12.669682 (2025).

29. Vazquez, R. et al. High frequency of CD95^+^ / CD45RA^−^ regulatory T cells defines an immunosuppressive profile associated with MDS progression. Br. J. Haematol. 208, 1993–2003 (2026).

30. Amir, E. D. et al. viSNE enables visualization of high dimensional single-cell data and reveals phenotypic heterogeneity of leukemia. Nat. Biotechnol. 31, 545–552 (2013).

31. Lê, S., Josse, J. & Husson, F. **FactoMineR** : An *R* Package for Multivariate Analysis. J. Stat. Softw. 25, (2008).

